# Relief of ParB autoinhibition by parS DNA catalysis and ParB recycling by CTP hydrolysis promote bacterial centromere assembly

**DOI:** 10.1101/2021.05.05.442573

**Authors:** Hammam Antar, Young-Min Soh, Stefano Zamuer, Florian P. Bock, Anna Anchimiuk, Paolo De Los Rios, Stephan Gruber

## Abstract

Three-component ParABS systems are widely distributed factors for plasmid partitioning and chromosome segregation in bacteria. ParB protein acts as an adaptor between the 16 bp centromeric *parS* DNA sequences and the DNA segregation ATPase ParA. It accumulates at high concentrations at and near a *parS* site by assembling a partition complex. ParB dimers form a DNA sliding clamp whose closure at *parS* requires CTP binding. The mechanism underlying ParB loading and the role of CTP hydrolysis however remain unclear. We show that CTP hydrolysis is dispensable for Smc recruitment to *parS* sites in *Bacillus subtilis* but is essential for chromosome segregation by ParABS in the absence of Smc. Our results suggest that CTP hydrolysis contributes to partition complex assembly via two mechanisms. It recycles off-target ParB clamps to allow for new attempts at *parS* targeting and it limits the extent of spreading from *parS* by promoting DNA unloading. We also propose a model for how *parS* DNA catalyzes ParB clamp closure involving a steric clash between ParB protomers binding to opposing *parS* half sites.

## Introduction

Faithful transmission of genetic material from one cell generation to the next is a prerequisite for survival and propagation. Chromosome segregation is spatially and temporally coordinated with DNA replication, cell division, and cell growth to maintain genome integrity and cell architecture over multiple generations. In bacteria, three-component ParABS systems promote faithful and efficient chromosome segregation (1-3). They are also important regulators of cell physiology with mutations leading to diverse and pleiotropic phenotypes in addition to chromosome partitioning errors. These phenotypes include defects in the control of DNA replication, chromosome organization, cell division, gene expression, motility, sporulation, and competence development (1, 4-14).

ParABS systems are present on most bacterial genomes and are also frequently encoded by low-copy number plasmids (15). They comprise 16 bp DNA sequence elements, called *parS* sites, the DNA-binding protein ParB, and the ATP-hydrolysing protein ParA. ParB proteins and *parS* sites together form the partition complex that serves as chromosome organizing centre (‘*parS* centromere’) (3, 16). Partition complexes are thought to follow ParA protein gradients on the bacterial chromosome thus becoming equidistantly positioned within the cell (12, 17, 18). They stimulate ATP hydrolysis by DNA-bound ParA dimers converting them into ParA monomers that dissociate from chromosomal DNA (5, 19-21). Using the same ‘diffusion-ratchet’ mechanism, ParABS is thought to promote plasmid partitioning (18, 22). The *parS* centromere also instructs another active DNA segregation mechanism, the Smc DNA loop extrusion motor. Starting from *parS* centromeres, the Smc complex aligns the two chromosome arms, helping to segregate nascent sister chromosomes (6, 23-26). ParABS furthermore regulates the initiation of DNA replication via the initiator protein DnaA in *B. subtilis*. Inability to convert ParA ATP-dimers into monomers (e.g. in a *ΔparB* mutant) leads to over-initiation of DNA replication (4) and as a consequence also blocks sporulation (27).

*parS* sequences are positioned near the replication origin, often in multiple copies scattered over a more or less wide replication origin region (<1 Mb) (15). ParB proteins accumulate in high numbers near a given *parS* site, leading to the formation of distinctive protein clusters in the cell (28-30). A related and essential feature of ParB is the ability to occupy not only the *parS* recognition sequence itself but also flanking DNA sequences. The spreading of ParB was first observed indirectly by its effects on plasmid supercoiling and silencing of *parS* proximal genes (31, 32). Chromosomal ParB is enriched in regions ranging from few kb up to ∼15 kb around *parS* sites in chromatin immunoprecipitation profiles (28, 33, 34). We and others have recently discovered that ParB spreading requires the binding of the unusual cofactor CTP by ParB (16, 35-37). Based on a nucleotide-ParB co-structure, we have proposed that ParB dimers form DNA sliding clamps that entrap chromosomal DNA in a *parS*-catalysed closure reaction to then slide onto *parS*-flanking DNA (37).

ParB proteins comprise three globular domains. The amino-terminal N domain forms a conserved binding pocket for the ribonucleotide CTP (36, 37). CTP binding (Kd ∼10 µM) (35, 37) is expected to be (nearly) saturated under standard physiological conditions (∼100-200 µM CTP). Upon contact with *parS* DNA, open ParB dimers convert into closed clamps by CTP-bound N domains forming interlocking dimers. N-gate closure is slow in the absence of *parS* DNA sequences. The *parS*-specific recognition of DNA originates from a helix-turn-helix (HTH) motif located in the ParB M domain (38). The carboxy-terminal C domain supports ligand-independent ParB dimerization as well as sequence-unspecific DNA binding (39, 40). CTP hydrolysis does not seem to be required for any step of ParB targeting and sliding *in vitro* (35, 37).

Here, we aimed to get a better understanding of the physiological role of CTP hydrolysis by generating ParB mutants in *B. subtilis*. We identified two CTP hydrolysis-defective mutants of ParB, which retained the ability to bind CTP, to close the N-gate, and to load onto *parS* DNA *in vitro*. We showed that CTP hydrolysis is essential for normal ParB focus formation *in vivo*. It serves two main functions by recovering CTP-locked ParB clamps trapped off the chromosomes and by restricting the extent of ParB spreading through DNA unloading. Despite the aberrant localization, CTP-hydrolysis-defective ParB mutants supported normal sporulation and Smc recruitment, but the mutant ParABS systems were unable to support chromosome segregation in the absence of Smc. Moreover, we present a model for *parS*-catalyzed ParB DNA loading based on *parS*-mediated relief of ParB autoinhibition.

## Results

Efficient chromosomal loading of ParB clamps requires the cofactor CTP and the catalyst *parS* DNA. CTP hydrolysis is thought to promote the reverse reaction, ParB clamp unloading. Here, we investigate the molecular mechanisms underlying these reactions as well as the physiological consequences of blocking CTP hydrolysis.

### Intra- and intermolecular tethering of N and M domains

In the crystal structure of a CDP-bound *B. subtilis* ParB dimer, the N domain of a given ParB chain closely associates with the M domain of the partner chain (Fig. 1A) (Fig. S1A) (PDB: 6SDK) (37). An equivalent domain-swap organization has recently been reported for a ParB paralogue, the *B. subtilis* protein Noc (PDB: 7NG0), and has previously been noted in the CTP-bound dimer of *Myxococcus xanthus* PadC (PDB: 6RYK) (36). This suggests that domain swapping is not a crystal packing artefact but a conserved feature of ParB and ParB-like proteins.

**Figure 1.**
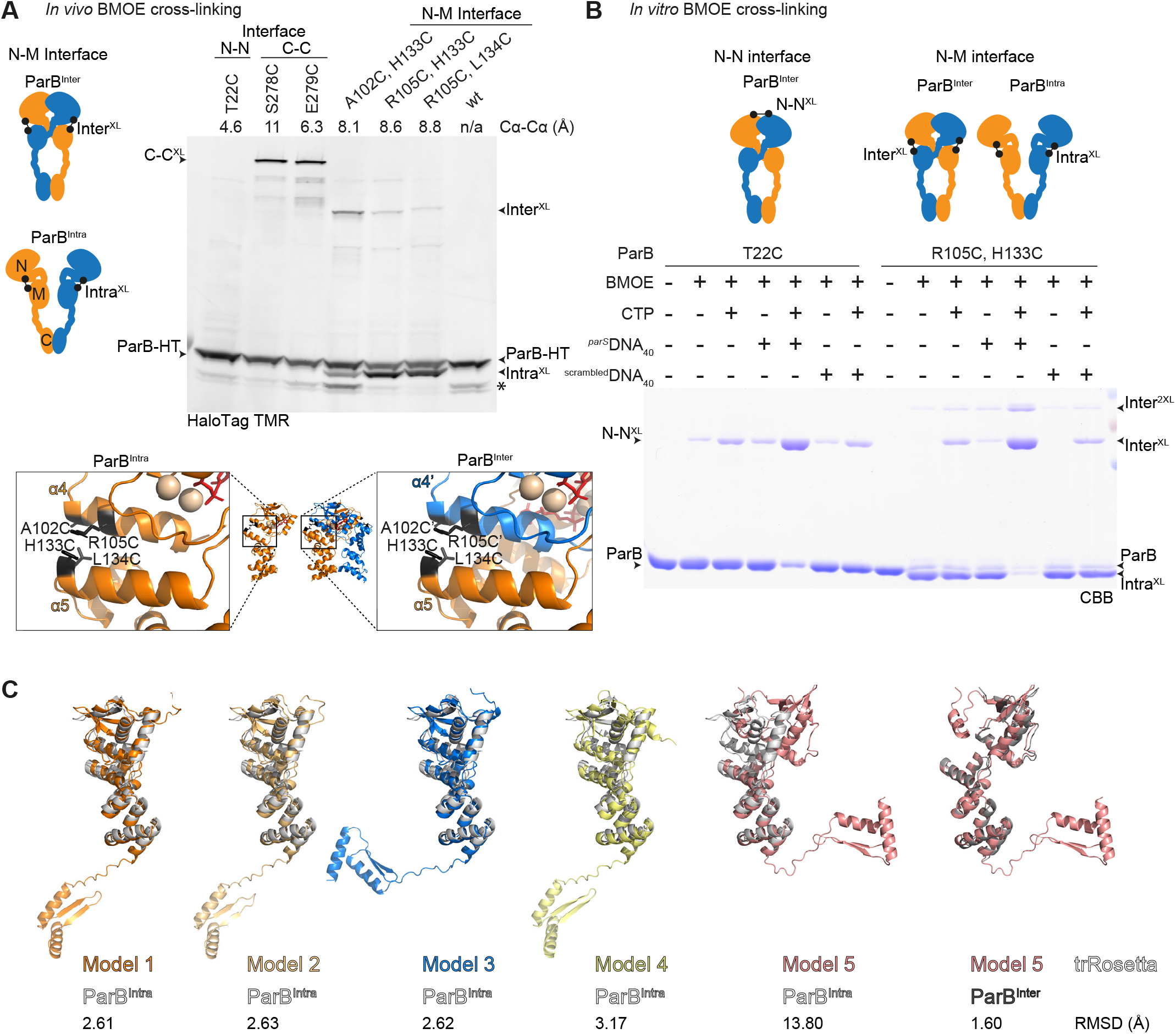
Intra- and intermolecular tethering of ParB domains. **(A)***In vivo* BMOE cysteine cross-linking with HaloTag (‘HT’) ParB variants. ParB-HT was labelled by addition of HaloTag-TMR ligand during cell lysis and cell extracts were analyzed by SDS-PAGE and in-gel fluorescence detection. Inter^XL^ denotes cross-links formed between the N domain of one protomer and the M domain of another. Intra^XL^ denotes cross-links between the N domain and the M domain of a given ParB protomer. Wild-type (‘WT’) ParB protein is naturally cysteine-free and failed to show any noticeable cross-linking. ParB dimers cross-linked at the C domain (C-C) ^XL^ were used as a positive control. Cα-Cα distances for corresponding pairs of cysteines on the ParB dimer structure (PDB: 6SDK) are denoted. The asterisk indicates a putative degradation product of ParB-HT. Structure inserts (bottom panels) denote the positions of the selected cysteine mutations on the ParB^Inter^ dimer structure (PDB: 6SDK) (right panel) and on a manually created ParB^Intra^ monomer model (left panel) (Fig. S2C). Of note, ChIP-qPCR assays verified that the T22C substitution did not adversely affect ParB localization on the chromosome (Fig. S7A). **(B)** *In vitro* BMOE cysteine cross-linking using cysteine pairs at the N-N interface (T22C) or the N-M interface. The cross-linking efficiency of ParB variants (at a monomer concentration of 10 μM) was tested with CTP (1 mM), 40 bp double-stranded *parS* DNA (^*parS*^DNA_40_) or equivalent DNA lacking *parS* sequences (^scrambled^ DNA_40_) (1 µM) in isolation or in combination. Bands were visualised on SDS-PAGE by Coomassie Brilliant Blue (CBB) staining. For quantification of cross-linking efficiencies see Fig. S2A. **(C)** ParB folding was predicted computationally by the *Robetta* server using default settings (*trRosetta*) with full-length *B. subtilis* ParB as input sequence (Uniprot: P26497). The top five search hits were then superimposed with the manually generated ParB^Intra^ model. Four superimpositions (1-4) displayed close structural resemblance. The remaining fold (5; left) showed reduced similarity with ParB^Intra^, but resulted in a good structural match when superimposed with the ParB^Inter^ structure (PDB: 6SDK) (5; right). Root mean square displacements (RMSD; in Å) are shown as an indicator of structural similarity.

To test if ParB adopts the domain-swap organization *in vivo*, we performed site-specific chemical cross-linking at the N-M interface using strains with a *halotag (HT)* fused to *parB* at the endogenous gene locus for the detection and quantification of cross-linked species (Fig. 1A) (37). Residues on helix α4 in the N domain (A102C and R105C) and on helix α5 in the M domain (H133C and L134C) were selected for cysteine mutagenesis (Fig. 1A). One of the four cysteine pairs (A102C/L134C) was omitted from the analysis as it exhibited a sporulation defect, being indicative of non-functional protein (42). The three other pairs did not grossly affect ParB function as indicated by the absence of a noticeable sporulation defect and by normal growth of the double cysteine mutants in a sensitized background harboring the *smc-pk3* allele (Fig. S1B) (24). If intermolecular N-M tethering does occur, cross-linking with the thiol-specific compound BMOE would generate a dimeric ParB-HT species that is expected to migrate more slowly through an SDS-PAGE gel. We indeed observed such a slowly migrating species (Inter^XL^) with all tested cysteine pairs (Fig. 1A) (Fig. S1C). The efficiency of intermolecular cross-linking was low and slightly higher only with one of the three cysteine pairs (A102C/H133C). Importantly, we also detected a more prominent, faster migrating cross-linked ParB-HT product, which we interpreted as a species (Intra^XL^) derived from intramolecular cross-linking of N and M domains (Fig. 1A).

This experiment suggested that N-M tethering occurs in two ways: within a protomer (‘ParB^Intra’^) and between two protomers (‘ParB^Inter^’) (Fig. 1, A and B). To elucidate how the domain-swap configuration is established, we next purified ParB(R105C, H133C) protein for *in vitro* cross-linking experiments. In the absence of ligands, BMOE cross-linking robustly produced the faster migrating species (Intra^XL^) and very little of two slowly migrating species (Inter^XL^ and Inter^2XL^), corresponding to single and double cross-linked ParB dimers (Fig. 1B) (Fig. S2A). We conclude that most or all ParB protein displayed the ParB^Intra^ configuration prior to ligand binding. Addition of CTP or ^*parS*^DNA_40_ alone did not alter the cross-linking pattern significantly, while addition of both strongly increased the abundance of the two dimeric species and in turn almost eliminated the intramolecular cross-linking. Similar results were obtained with ParB(A102C, H133C) (Fig. S2B). The requirement for both ligands mimicked what was observed for N-gate closure by T22C cross-linking (Fig. 1B) (37). N-gate closure thus goes along with the conversion of ParB^Intra^ into ParB^Inter^.

### A mechanism for *parS*-catalyzed ParB loading based on ParB autoinhibition

The closed ParB^Inter^ state is thermodynamically favorable in the presence of CTP (and CTPγS but not CDP)(37). The conversion of CTP-bound ParB^Intra^ to ParB^Inter^ however is slow without *parS* DNA (i.e. a kinetically inhibited reaction) (Fig. S2C) (37). What inhibits the reaction in the absence of *parS* DNA is unclear. Disengagement of intramolecular N-M tethers in both ParB^Intra^ protomers is presumably a prerequisite for forming the closed N-gate built from two ParB^Inter^ protomers. We thus wondered whether the intramolecular N-M tether inhibits N-gate closure. If so, ParB^Intra^ would correspond to an autoinhibited form of ParB. Stochastic N-M disengagement is likely inefficient due to rapid reengagement of the juxtaposed N and M domains, even in case of limited N-M tether stability.

To uncover how such ParB autoinhibition might be overcome by *parS* DNA binding, we generated a structural model of ParB^Intra^ by manually reconnecting the chains at the N to M junction of the CDP-bound ParB dimer (PDB: 6SDK) (Fig. S3A). We also computationally predicted ParB structure (de novo) using trRosetta (43). One of the top five hits closely resembled the ParB^Inter^ structure, while the other four matched well with the manually created ParB^Intra^ model, providing unbiased and independent support for the existence of these states (Fig. 1C). Next, we superimposed two ParB^Intra^ chains with opposing halves of a *parS* site (PDB: 4UMK), using the helix-turn-helix motif in the M domain as guide for structural alignment. We found that a single ParB^Intra^ protomer can readily accommodate a *parS* half site, but two ParB^Intra^ protomers clash with one another when bound to opposing half sites, indicating that a ParB^Intra^ dimer fails to strongly bind *parS* DNA (Fig. 2A). However, when the N domain was manually detached from the M domain (ParB^Untethered^) in at least one of the two protomers, then binding of both HTH motifs to the half sites of a given *parS* DNA sequence becomes feasible without steric clash (Fig. 2A). Thus, we propose that *parS* DNA catalyzes N-gate closure by preventing N-M reengagement and selecting and stabilizing a ParB^Untethered^ state (Fig. 2B). With one protomer being kept in this—otherwise thermodynamically unfavorable—state, N-gate closure may proceed efficiently as soon as the other protomer stochastically happens to adopt the same state. Consistent with this hypothesis, the N domain of at least one of the two *parS*-bound ParB chains has been observed in a partially unfolded state in available ParB-*parS* co-crystal structures (PDB: 4UMK) (38) (PDB: 6T1F, personal communication with Tung Le). Accordingly, open ParB dimers exist as an ensemble of states with protomers in ParB^Intra^ and ParB^Untethered^ conformations, while closed ParB dimers are built from ParB^Inter^ protomers.

**Figure 2.**
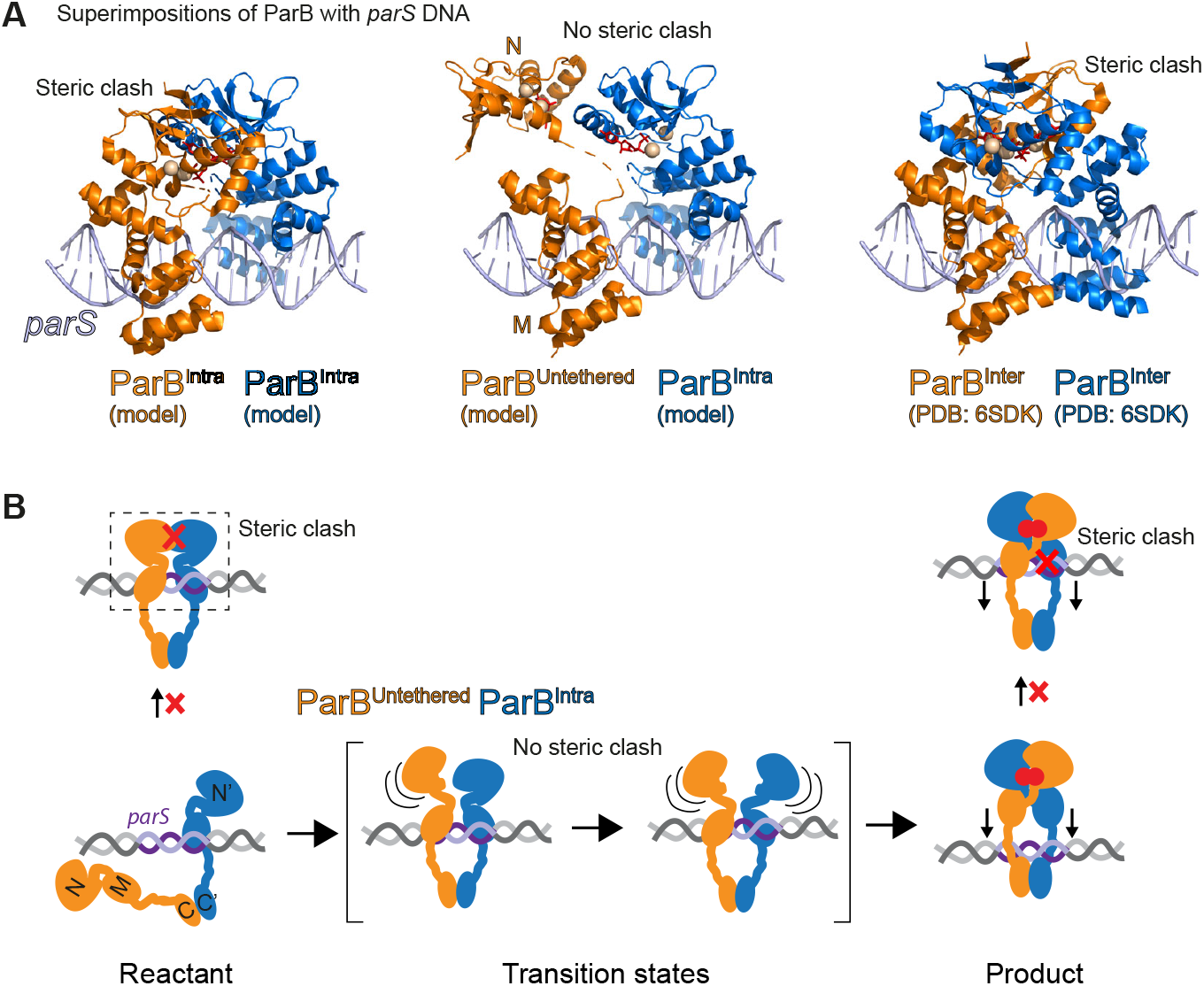
Steric hindrance of *parS* DNA binding by ParB^Intra^ dimers and ParB^Inter^ dimers. **(A)** Superimposition of ParB chains with *parS* DNA. Structural models of the ParB^Intra^ state were manually created from the ParB^Inter^ structure (PDB: 6SDK) (Fig. S3A) or computationally predicted (See Fig. 1C). Binding to *parS* DNA was modelled by superimposition of ParB^Intra^ (left panel) and ParB^Inter^ with a *H. pylori* ParB-*parS* co-crystal structure (PDB: 4UMK) using the HTH motif as a guide for structural alignment yielding a steric clash of the N domains. A ParB^Untethered^ model was generated by manually detaching the N domain from the M domain. Superimposition with *parS* DNA (middle panel) was possible without steric hindrance. **(B)** A model for *parS*-catalysed N-gate closure. ParB dimers fail to bind to both *parS* half sites simultaneously due to steric clash. Only when the N domain is partially unfolded or untethered from the M domain in at least one ParB protomer, does the binding to both *parS* half sites become feasible (‘transition state’). The interlocking of the two ParB protomers (ParB^Inter^ N-M) re-generates the steric clash with *parS* DNA binding, thus releasing the closed ParB dimer from *parS* DNA.

A prediction from this Brownian-ratchet reaction scheme is that the steric clash between two ParB^Intra^ protomers at *parS* DNA is crucial for catalysis. To artificially eliminate the steric clash, we inserted an increasing number of spacer nucleotides (adding 1 to 8 bp) between the two *parS* half sites in ^*parS*^DNA _40_. We then tested for the ability of the modified *parS* DNA to stimulate CTP hydrolysis—an indirect readout for the efficiency of N-gate closure. We found that all altered *parS* sites failed to stimulate CTP hydrolysis. Intriguingly, DNA molecules with intermediate spacer lengths (2-5 bp) also inhibited the reaction when present at stoichiometric amounts (Fig. S3B), presumably by allowing unhindered ParB^Intra^ binding to both *parS* half sites. These observations are consistent with the idea of a steric clash between *parS*-bound ParB^Intra^ protomers being the basis of *parS*-catalyzed ParB DNA loading. Notably, ParB binding to isolated *parS* half sites has been observed in ChIP-Seq profiles for *Pseudomonas aeruginosa* ParB, particularly upon ParB over-expression (44), and possibly also in other organisms (45).

### CTP hydrolysis-defective ParB mutants

We next focused on the reverse reaction, i.e. ParB clamp opening, which is presumably supported by CTP hydrolysis. The ParB CTPase is a recently discovered member of a larger family of proteins with diverse enzymatic activities and cofactors (37, 46). Little is yet known about the mechanism of ParB CTP hydrolysis. In ATP and GTP hydrolysis, a water molecule attacks the γ-phosphate moiety to hydrolyze the scissile bond between β- and γ-phosphates. The water molecule is usually activated for nucleophilic attack by an acidic residue, as for example in the Walker B motif. To identify residues with equivalent functions in ParB, we biochemically characterized ParB proteins harboring mutations in selected active site residues. The desired mutants were expected to be defective in CTP hydrolysis but retain their ability to bind CTP, engage the N domains, and entrap *parS* DNA. The ParB-CDP co-crystal structure (PDB: 6SDK) highlighted two glutamate residues (at position 78 and 111) at the CTP binding pocket (Fig. 3A) (37). These residues belong to widely conserved sequence motifs (GE_78_RRY/F and E_111_NLQR) but they are absent from *M. xanthus* PadC protein which binds CTP but does not hydrolyze it (with F348 in PadC corresponding to *B. subtilis* ParB E78 and PadC lacking the ENLQR motif altogether) (Fig. S5A) (36). Examining the PadC-CTP interaction map confirmed that residue F348 is suitably positioned next to the CTP γ-phosphate (Fig. S5B). The glutamate residues were chosen as candidates for further analysis and replaced separately by glutamine (Q). Substitutions for alanine or histidine produced

**Figure 3.**
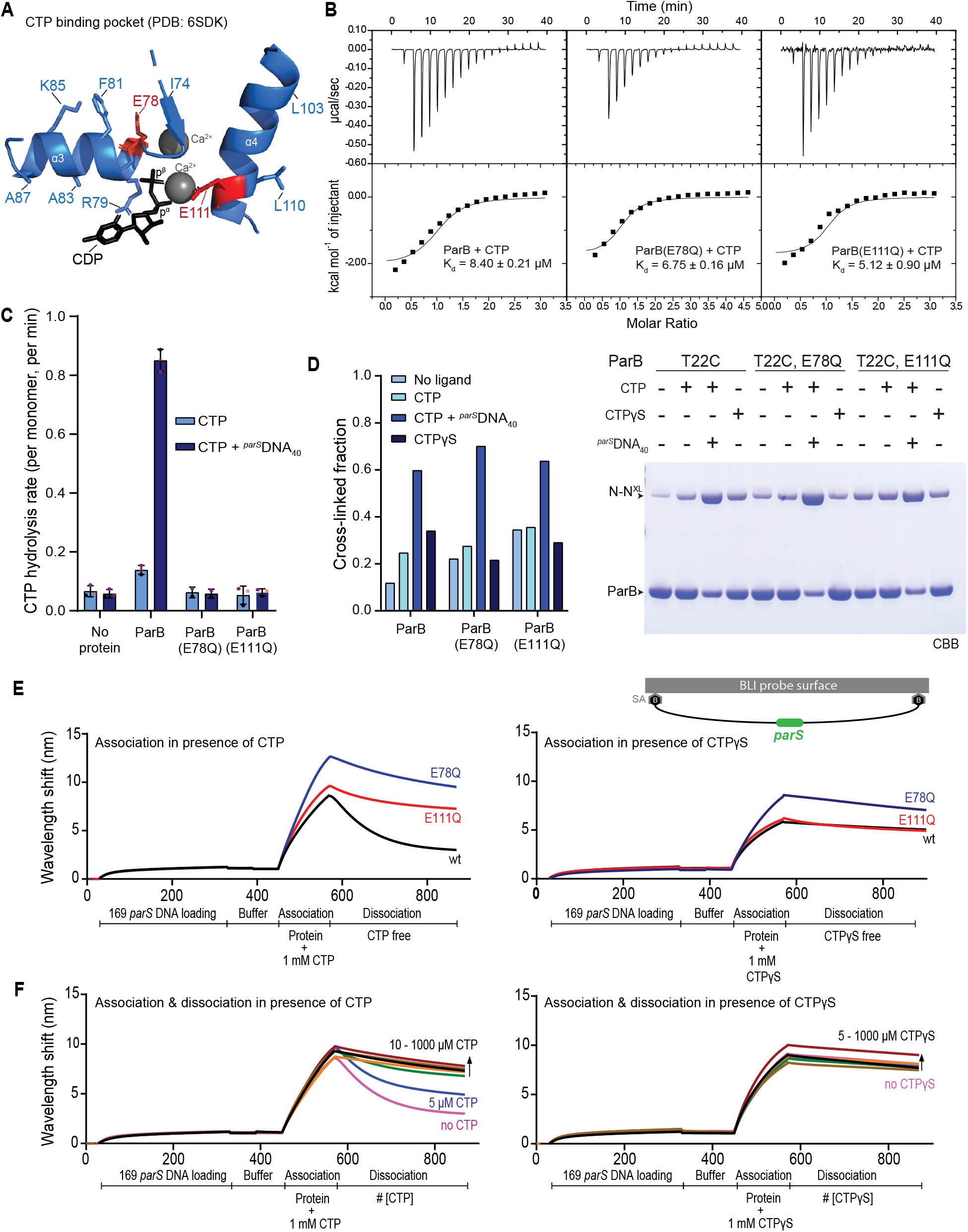
Hydrolysis-defective mutants of ParB. **(A)** The CTP binding pocket found in a ParB-CDP co-crystal structure (PDB: 6SDK) in cartoon representation. Conserved residues are shown in stick representation. Candidate catalytic residues, glutamate at position 78 and 111, are highlighted in red colours. **(B)** ParB-CTP affinity measurements by ITC. Wild-type or mutant ParB, at a monomer concentration of 80-120 μM, was injected with 2 mM CTP solution. The K_d_ from a typical experiment is given. For details see Materials and Methods. **(C)** Rate of CTP hydrolysis assayed by colorimetric measurement of free inorganic phosphate (Malachite green assay). 10 µM of ParB was incubated with 1 mM CTP with or without 1 µM ^*parS*^DNA_40_. Mean values and standard deviation from three repeat measurements are reported. Individual data points are shown as dots. **(D)** ParB N-gate closure measured by *in vitro* BMOE cross-linking of ParB(T22C). Final reactions contained 10 µM ParB with or without 1 mM NTP and 1 µM ^*parS*^DNA_40_ in Mg2+ containing buffer. N-N^XL^ denotes cross-linked species. Cross-linked species were visualised on SDS-PAGE and staining with Coomassie Brilliant Blue (CBB) (right panel). Quantification of bands (left panel) was performed with ImageQuant (GE Healthcare). **(E)** ParB loading on immobilized *parS* DNA measured by biolayer interferometry (BLI). Interaction measured between 1 µM of ParB variant protein and immobilized biotin-labelled 169 bp *parS* DNA, in Mg2+ containing buffer, with 1 mM CTP and CTPγS (left and right panel, respectively). For dissociation, an equivalent buffer lacking ParB protein and nucleotide was used. **(F)** Dissociation of wild-type ParB form *parS* DNA in the presence of nucleotide. Same as in (E) but with increasing concentrations of CTP (left panel) and CTPγS (right panel) in the dissociation buffer.

similar initial results but were deemed more intrusive and thus omitted from further investigations (Fig. S7C). The E78Q and E111Q mutant proteins were recombinantly expressed and purified. We assayed for CTP binding affinity, for the rate of CTP hydrolysis, for the efficiency of N-gate closure, and for ParB loading onto *parS* DNA. Our results showed that the E78Q and E111Q mutants failed to display appreciable levels of CTP hydrolysis as measured by the release of inorganic phosphate, while a wild-type control protein hydrolyzed CTP with a basal rate of approximately 0.15 min^-1^, which was stimulated by the presence of *parS* DNA to about 1 min^-1^, comparable with published results (Fig. 3C) (37). All three proteins bound CTP with similar affinity (Kd ∼5-8 µM) as judged by ITC (Fig. 3B), and they had roughly comparable efficiencies of N-gate closure with CTP and *parS* DNA as measured by T22C cross-linking (Fig. 3D). They also loaded onto *parS* DNA in the presence of CTP based on biolayer interferometry (BLI) (Fig. 3E). In brief, BLI allowed for the immobilization of a double biotin labeled 169 bp *parS* DNA fragment on a streptavidin-coated biosensor tip (35). Binding of ligands, including ParB protein, is inferred from a wavelength shift in the reflected light. Association with *parS* DNA appeared normal or even slightly improved with the EQ mutants in the presence of CTP or the non-hydrolysable analog CTPγS. When shifting the biosensor from the CTP-containing loading buffer to a CTP-free dissociation buffer, wild-type ParB protein was released from DNA with an estimated apparent rate of about 0.4 min^-1^, while the two EQ mutants displayed a significantly longer residence time on DNA. These differences in the dissociation rates between wild-type and EQ proteins were eliminated when CTPγS was used instead of CTP during loading. Of note, we observed that the presence of CTP in the dissociation buffer significantly reduced the off-rate of wild-type ParB (Fig. 3F), suggesting that *B. subtilis* ParB can efficiently exchange CDP for CTP without dissociating from DNA as previously reported for *C. crescentus* ParB (35). We determined a CTP concentration for half-maximal ParB retention of about 5-10 µM, which is in agreement with the CTP affinity measured by ITC (Fig. 3F).

CTPγS promotes slow but robust N-gate closure even in the absence of *parS* DNA (37). CTP is unable to do so in wild-type ParB protein, presumably due to its hydrolysis to CDP with ParB^Intra^ being the most populated CDP-state (and apo-state). As expected, we found that the EQ mutants supported N-gate closure equally well with CTP and CTPγS (Fig. S4), thus providing further support for the notion that the EQ mutants are defective in CTP hydrolysis and that CTP hydrolysis counteracts N-gate closure. Notably, the N-gate closure reaction was somewhat slower in E78Q and more so in E111Q when compared to wild type, possibly implying that the mutant proteins are either slightly more strongly autoinhibited (more stable ParB^Intra^) or less stably closed (less stable ParB^Inter^) (Fig. S4). Altogether, we conclude that the EQ mutants are defective in CTP hydrolysis but support all other biochemical functions normally or near normally *in vitro*.

### CTP hydrolysis is dispensable for SMC recruitment but not for ParABS function

To elucidate the physiological consequences of defective CTP hydrolysis, we transferred the ParB(EQ) mutations into *B. subtilis* by allelic replacement. We observed that neither the E78Q nor the E111Q mutation resulted in a noticeable sporulation defect as judged after extended periods of incubation on nutrient-rich agar plates (Fig. 4A), implying that the regulation of DNA replication is only mildly or not at all perturbed in the mutants in contrast to the *parB* in-frame deletion mutant (*ΔparB*). Next, we combined the EQ mutations with a hypomorphic allele of the *smc* gene, *smc-pk3*, to sensitize cells for defects in chromosome organization and segregation (24). Again, unlike the *ΔparB* mutant, the EQ mutants supported robust growth of the *smc-pk3* strain (Fig. 4B), demonstrating that the *parB(EQ)* mutants supported chromosome segregation well, presumably by promoting the loading of Smc-ScpAB onto the chromosome. To test this more directly, we next performed chromatin immunoprecipitation (ChIP) using antiserum raised against the *B. subtilis* Smc protein. ChIP-Seq showed that the chromosomal distribution of the Smc protein was similar in the EQ mutants and wild type, and distinct from the *ΔparB* mutant (Fig. 4C). ChIP-qPCR analysis of the same samples suggested that the levels of enrichment of the Smc protein were slightly reduced at the *parS-359* site and the replication origin (‘*dnaA*’) in the EQ mutants, but clearly not as much reduced as in *ΔparB*, together providing further support for the notion that CTP hydrolysis is largely dispensable for Smc recruitment and loading (Fig. S6B).

**Figure 4.**
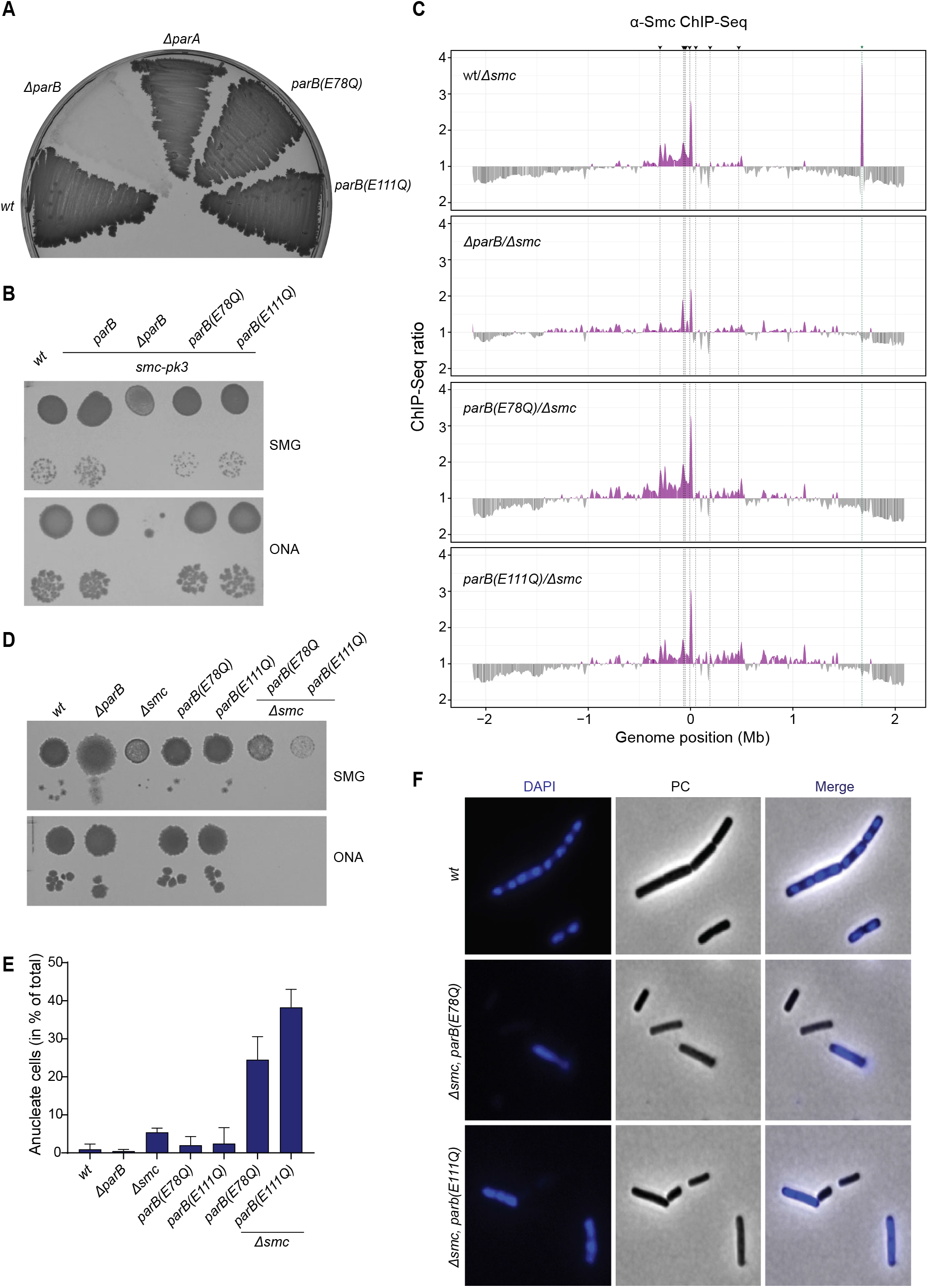
Cellular phenotypes of ParB CTP hydrolysis mutants. **(A)** Sporulation of strains with indicated *parB* alleles. Image taken after 3 days of incubation on nutrient rich agar plate at 37°C. **(B)** Growth assay by dilution spotting of wild-type *B. subtilis* and strains carrying *parB* alleles in an *smc-pk3* background (hypomorphic mutant). 9^2^ and 9^5^-dilutions were spotted on minimal medium agar plates (SMG) and rich medium agar plates (ONA) and imaged after 24 and 16 hours, respectively. **(C)** Chromatin-immunoprecipitation coupled to deep sequencing (ChIP-Seq) using α-Smc serum. Panels show ratio plots of Smc enrichment against that of *Δsmc*. All ChIP-Seq profiles were split into 1 kb bins with the origin of replication placed at position 0 Mb. The ratio was calculated by dividing the higher value by the lower, if the *condition/Δsmc* ratio was > 1, it was plotted above the genome position axis (purple colour). If the *Δsmc/condition* ratio was > 1 the ratio was plotted below the axis (grey colour). Black dashed lines and arrows represent the 8 prominent *parS* sites. Of note, the peak at position +1.8 Mb (green dashed line) marked by asterisk represents enrichment of the *smc* gene presumed to be a contamination. **(D)** Growth assay by dilution spotting of wild type *B. subtilis* and strains carrying parB alleles in a smc deletion background (‘*Δsmc*’). 9^2^ and 9^5^-fold dilutions were spotted on minimal (SMG) and rich medium (ONA) and imaged after 24 and 16 hours, respectively. **(E)** Quantification of the fraction of anucleate cells using ImageJ with the microbeJ plugin (Ducret et., al 2016) in *parB* and *parB, Δsmc* mutants. Five fields of view were captured randomly for each strain containing roughly 100 cells per field of view. All cells in the field of view were counted (unless they’re too close to the border of the field of view). Cell outline and focus detection were reviewed manually to ensure accurate segmentation and true focus (maxima) detection. The percentage of anucleate cells is reported as the number of cells without maxima divided by the total cell count and corrected to 100. Mean value of anucleate cells per field of view is reported for each strain with the standard deviation.**(F)** Example images showing DAPI DNA staining (in blue colours) for wild type, *Δsmc parB(E78Q)*, and *Δsmc parB(E111Q)* strains of *B. subtilis*.

Next, we investigated the role of ParB CTP hydrolysis in the core function of the ParABS system. To completely rule out effects of ParB-Smc interactions on chromosome organization and segregation, we constructed the *parB(EQ)* mutants in a *Δsmc* strain. We note that the isolation of such double mutant strains was difficult, similar to the mutant of both *Δsmc* and *ΔparB*. The resulting strains failed to grow on nutrient-rich medium and also displayed very poor growth characteristics under nutrient limiting conditions and at reduced temperatures, thus being markedly sicker than the *Δsmc* strain (Fig. 4D). Further investigation revealed that the two double mutants (*parB(EQ), Δsmc*) showed strong chromosome segregation defects with around 25 % and 35 % of cells lacking DNA staining (‘anucleate’) in E78Q and E111Q, respectively (Fig. 4, E and F). We conclude that ParB CTP hydrolysis becomes critical for efficient chromosome segregation in the absence of Smc function in *B. subtilis*, strongly suggesting that proper function of ParABS in chromosome segregation critically relies on ParB CTP hydrolysis.

### CTP hydrolysis prevents excessive ParB spreading and off-target accumulation of closed ParB

To reveal how CTP hydrolysis may promote ParABS function while being apparently dispensable for Smc recruitment, we studied the cellular localization and chromosomal distribution of ParB protein in *B. subtilis*. We performed ChIP-qPCR using antiserum raised against full-length *B. subtilis* ParB protein. The enrichment of ParB(E78Q) was clearly reduced at the *parS-359* site and the neighboring gene *parA* (*soj*) (Fig. 5A). This reduction was even more pronounced for ParB(E111Q) (Fig. 5A). Deep sequencing of the ChIP eluates resulted in reduced read counts in the EQ mutants at *parS*-359 and at other *parS* sites, again with the E111Q variant displaying a more drastic phenotype. Moreover, the shape of the ParB distribution at *parS* sites was clearly altered in both EQ mutants. The enriched region was significantly expanded by extending further onto one side of *parS*—away from the replication origin—but not onto the other (Fig. 5B). The width of the peak at *parS-359* increased from approximately 15 kb for wild type to roughly 40 kb for the two EQ mutants. The peak position was also slightly shifted away from the *parS* site in the direction of the extended shoulder. This suggested that the ParB(EQ) mutants exhibited excessive spreading, presumably by an increased chromosome residence time in the absence of CTP hydrolysis. This excessive spreading of ParB(EQ) was highly asymmetric, putatively due to the impediment of spreading in one direction by other chromosomal processes such as head-on transcription or DNA replication. If so, then CTP hydrolysis might reduce the frequency or impact of such encounters in wild-type cells. How this might be accomplished is unclear.

**Figure 5.**
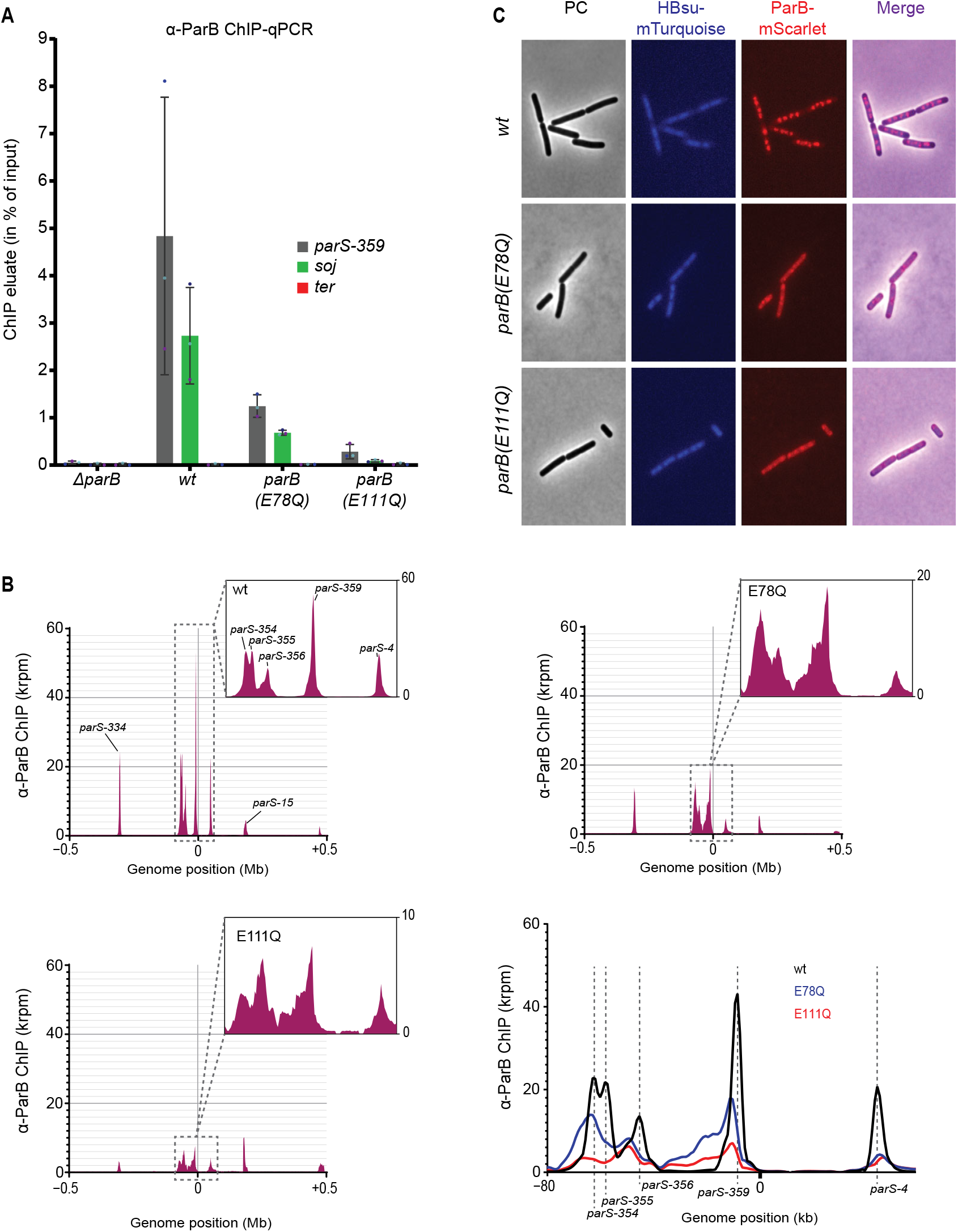
Localization defects of CTP hydrolysis-defective ParB mutants. **(A)** Chromatin-immunoprecipitation coupled to quantitative PCR (ChIP-qPCR) using α-ParB serum. Mean values and standard deviation from three biological replicates are reported. Individual data points are shown as dots. Enrichment of ParB was tested at the three loci: *parS-359, parA (soj)*, and the terminus (*yocGH*). **(B)** Chromatin-immunoprecipitation coupled to deep sequencing (ChIP-Seq) using α-ParB serum. Top left panel shows the sequence read distribution profile for wild-type ParB in a region surrounding the replication origin, top right and bottom left panels show the profiles for *parB(E78Q)* and *parB(E111Q)*, respectively. Note the different scales of the Y-axis in the zoomed-in windows. Bottom right panel shows an overlay of the three profiles in a region including the *parS-354* and *parS-4* sites. All ChIP-Seq profiles were split into 1 kb bins with the origin of replication placed at position 0 Mb. Plots in bottom right panel have been smoothened for presentation. **(C)** Fluorescence microscopy of *B. subtilis* with different alleles of *mScarlet*-tagged *parB* (in red colours) in combination with *mTurquoise*-tagged *hbs* (in blue colours).

We then investigated the cellular localization of wild-type and mutant ParB-mScarlet fusion proteins by live-cell imaging (Fig. 5C). These strains also expressed a *mTurquoise*-tagged ectopic copy of *hbs*, encoding for an abundant nucleoid associated protein with sequence-unspecific DNA-binding properties. Wild-type ParB-mScarlet formed the characteristically bright foci that are known to colocalize with *parS* sites. In contrast, the two hydrolysis mutants displayed more diffusive signal with only faint foci being detected. Of note, the mScarlet fusion mildly reduced the enrichment of ParB at two tested loci as judged by ChIP-qPCR (Fig. S7B).

In summary, we observed defects in the accumulation and the distribution of ParB(EQ) proteins near *parS* sites. The total level of protein in ParB-mScarlet foci was decreased in the two EQ mutants, mildly in E78Q and strongly in E111Q. The occupancy of the *parS*-proximal regions was also reduced, while the occupancy of more distal positions on one of the two *parS*-flanking regions was increased.

### CTP hydrolysis-defective ParB mutants accumulate in an alternate state on the chromosome

To elucidate the state of ParB(EQ) proteins in the cell, we next employed *in vivo* cysteine cross-linking using ParB-HT fusion proteins. Presumably due to CTP hydrolysis dominating over N-gate closure *in vivo*, T22C cross-linking of otherwise wild-type ParB-HT protein was barely detectable (Fig. 1A, 6A), as reported previously (37). Cross-linking of R105C/H133C also showed mostly the ParB^Intra^ configuration (Fig. 1A, 6A). The EQ mutants, however, robustly generated ParB N-N cross-links with T22C and ParB^Inter^ N-M cross-links with R105C/H133C (Fig. 6A). Thus, in contrast to wild-type ParB, the EQ mutants accumulated predominantly in the closed form *in vivo*. This finding provided further support for the notion of defective CTP hydrolysis in the EQ proteins.

**Figure 6.**
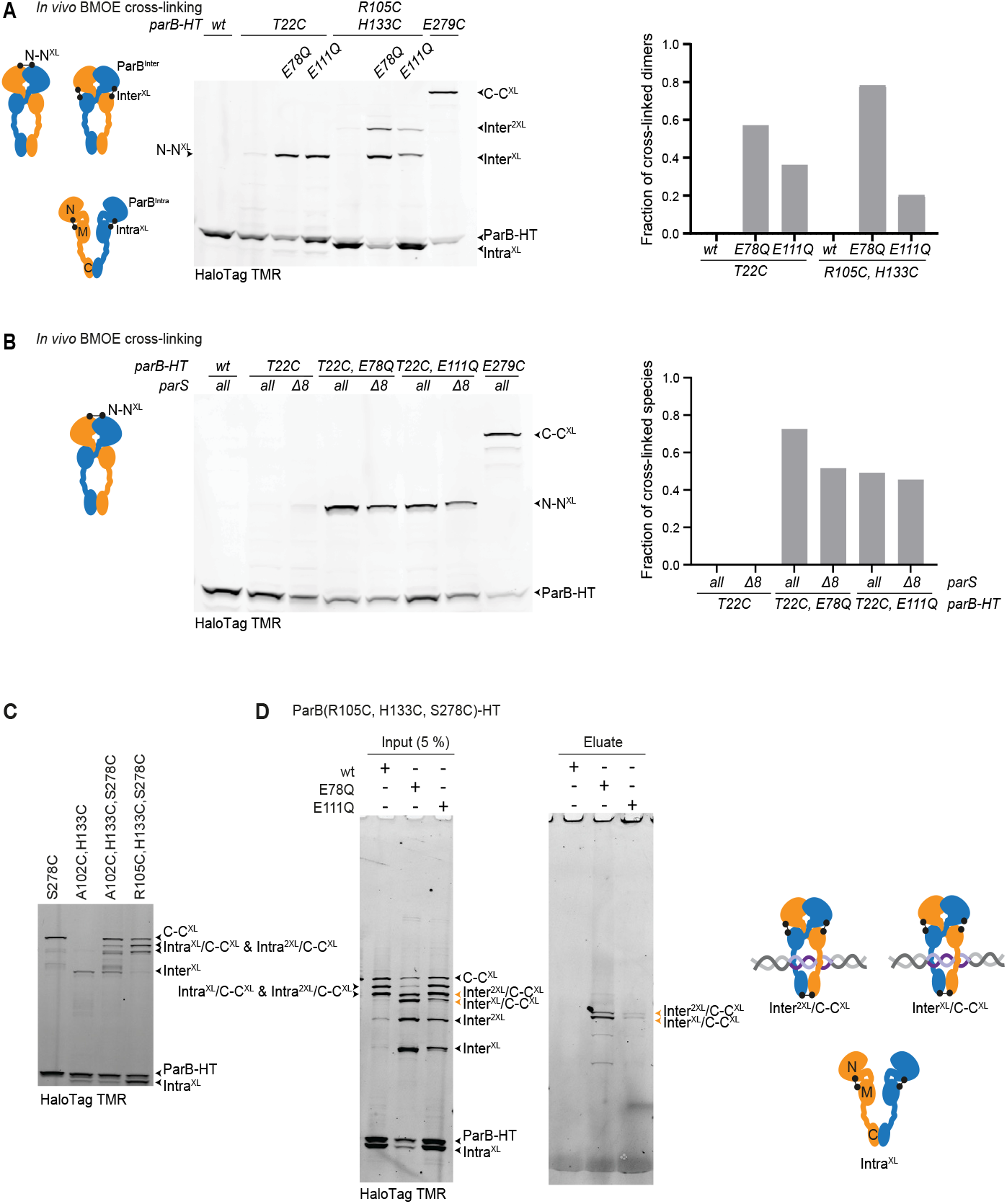
Structural states of wild-type ParB and ParB(EQ) *in vivo*. **(A)** *In vivo* BMOE cysteine cross-linking assay with ParB-HT variants. Cross-linked cell fractions were lysed and analysed by SDS-PAGE and in-gel fluorescence detection of ParB-HT species (left panel). Cross-linking efficiencies were calculated by quantifying band intensities of monomeric species of ParB (ParB-HT for T22C; ParB-HT and Intra^XL^ for R105C/H133C) and cross-linked dimeric species (N-N^XL^ for T22C; Inter^XL^ and Inter^2XL^ for R105C/H133C) (right panel). Residue E279C in domain C was used a positive control for cross-linking. **(B)** Cysteine cross-linking in strains lacking *parS* sites. As in (A) using strains lacking the 8 *parS* sites. **(C)** Cysteine cross-linking in strains with multiple cysteine pairs in ParB-HT. As in (A) for identification of cross-linked species used in (D). Note that species with single and double Intra^XL^ as well as C-C^XL^ cannot be unambiguously identified. Bands corresponding to circular species (as seen in panel D) are not clearly discernible in the absence of hydrolysis mutations. **(D)** Chromosome entrapment assay using strains harbouring ParB(R105C, H133C, S278C)-HT in combination with E78Q or E111Q. Input and output samples were analysed by SDS-PAGE and in-gel fluorescence detection. With E78Q and E111Q, differing levels of cross-linked ParB species were found in the eluate samples. Covalent circular species of ParB-HT are denoted by arrowheads in orange colours. Of note, CTP hydrolysis-proficient ParB did not generate covalent circular species in the input samples (see also panel C). Cross-link reversal leads to the generation of minor amounts of non-circular species in the eluates (48-50).

The fact that EQ proteins showed more robust N-gate closure compared to wild type (Fig. 6A) implies that a large fraction of wild-type ParB in partition complexes harbors an opened N-gate. Moreover, the poor localization of the EQ mutants (and in particular of E111Q) to partition complexes indicated that they may have undergone N-gate closure without *parS* stimulation or have dissociated from *parS* DNA after N-gate closure. To discriminate between these two possibilities, we have repeated the T22C cross-linking experiment in a strain lacking 8 strong *parS* sites (47). We found that T22C cross-linking was only mildly reduced in both EQ proteins (Fig. 6B), suggesting that a significant fraction of closed ParB clamps are formed without *parS* stimulation. CTP hydrolysis likely recovers such futile clamps in wild-type ParB.

As a measure for chromosome association irrespective of chromosome localization, we finally performed chromosome entrapment assays. Briefly, we used cysteine cross-linking to generate covalently closed circular species of ParB-HT *in vivo* (Fig. 6C) to then detect their co-isolation with chromosomal DNA in agarose beads under protein denaturing conditions (48, 49). Only cross-linked species that entrap the chromosomal DNA double helix are retained in the agarose matrix. Using R105C/ H133C at the N-M interface in combination with S278C at the C-C interface, we found that significant levels of cross-linked ParB(E78Q) species were retained in agarose beads (Fig. 6D). The retention of ParB(E111Q) species was comparatively low. We conclude that a considerable fraction of ParB(E78Q) entraps chromosomal DNA as a CTP-closed clamp, presumably partly near *parS* and partly distal from *parS*. Most of the ParB(E111Q) [and some of ParB(E78Q)], however, appeared to form closed ParB dimers without entrapping chromosomal DNA. Of note, as wild-type equivalents do not show robust ParB^Inter^ cross-linking with R105C/H133C, no information on hydrolysis-proficient ParB was obtained in this experiment. Together our results indicate that ParB and ParB(EQ) occupy distinct states in the cell and when loaded onto the chromosome, which may explain at least in part the defects in ParABS function in the EQ mutants. A model figure summarizes these findings (Fig. 7A).

**Figure 7.**
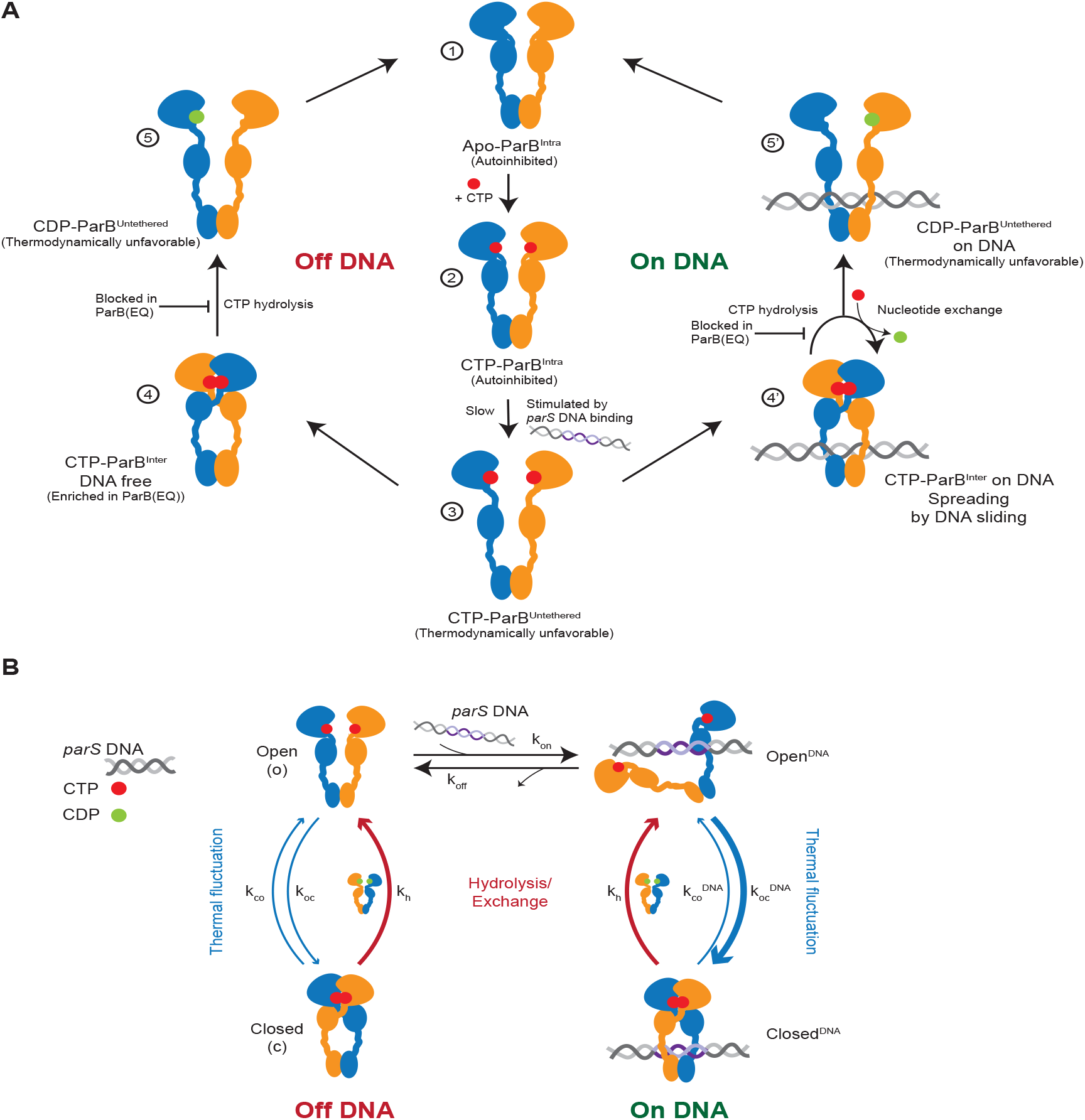
Model for partition complex assembly by the ParB CTPase. **(A)** A model for ParB CTP hydrolysis and *parS* DNA catalysis in partition complex assembly. CTP binding by ParB^Intra^ (1) and (2). The N domain spontaneously dissociates from the M domain to generate ParB^Untethered^ (3). This transition is thermodynamically unfavorable and slow but promoted by *parS* DNA binding (for details see Fig. 2B). Subsequent clamp closure is rapid off DNA (left side) (4) and on *parS* DNA (right side) (4’). Due to a steric clash between *parS* DNA and the two M domains (Fig. 2B), the DNA is released from *parS* DNA and located in a lumen formed by the M and C domains (4’). ParB spreads away from *parS* DNA by 1D diffusion onto *parS* flanking DNA. CTP hydrolysis by DNA-bound ParB leads to nucleotide exchange or to unloading from DNA (5’). After unloading, reengagement of the N and M domains within a given ParB protomer regenerates the autoinhibited open ParB dimer (1). The partially saturated nucleotide binding sites (5 and 5’) indicate that nucleotide release and exchange may happen before or after transition to 5 and 5’. Clamp opening upon hydrolysis limits excessive spreading of ParB by unloading from DNA (4’->5’) and allows for the recycling of DNA-free ParB clamps (4->5). The absence of CTP hydrolysis in ParB(EQ) mutants leads to the excessive ParB spreading by long-term chromosome residence (4’) as well as to the accumulation of DNA-free ParB clamps (4). **(B)** A simplified reaction scheme for CTP-hydrolysis promoted enrichment of ParB at *parS* sites. Under the assumption of fast nucleotide exchange, only the CTP-bound states are considered. DNA binding and unbinding from the closed form is expected to be slow and neglected in the simple model.

### CTP hydrolysis promotes ParB/*parS* ultra-affinity

The detailed model (Fig. 7A) could be further coarse grained to a simple scheme for *parS* DNA association (Fig. 7B) by means of a few reasonable approximations. Only the open form of ParB can engage the substrate, whereas closed ParB cannot bind or release DNA substrates, because steric clashes make these processes extremely unlikely, if not impossible. Furthermore, ParB without nucleotides is a transient, poorly populated state, as could be inferred from the longer persistence of already bound ParB on DNA in the presence of CTP in BLI experiments (Fig. 3F). Thus, we captured the transition from CDP-bound to CTP-bound ParB by means of a simple transition. The resulting model can be analytically solved (see Materials and Methods), giving an estimate of the “apparent”, observed, dissociation constant 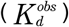 of ParB for DNA:

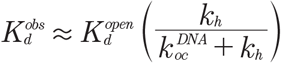

The *parS* sequence greatly accelerates ParB closure 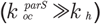 thus decreasing the value of the observed dissociation constant. In the case of non-specific DNA, however, the rate of ParB closure is unaffected and likely slower than the hydrolysis rate, resulting in a dissociation constant which is the same as the one of open ParB. The same observed dissociation constant also applies to hydrolysis deficient mutants at equilibrium (see Materials and Methods).

## Discussion

The partition complex is critical for ParABS function. The assembly has been intensely studied over several decades. The recently discovered requirement for CTP binding by ParB in partition complex assembly opened new avenues for investigation. Here, we focused on the mechanistic basis of *parS* DNA catalysis of ParB clamp loading and on the physiological role of ParB CTP hydrolysis in partition complex assembly and chromosome segregation.

### *parS* DNA: A B-form DNA double helix as catalyst

Ribozymes are naturally occurring nucleic acid catalysts that generally utilize folded single-stranded RNA to catalyze a variety of chemical reactions. Unlike ribozymes, *parS* DNA appears to work as catalyst in the standard B-form of double-stranded DNA. Based on structural information and chemical cross-linking, we here propose a molecular mechanism for *parS* catalysis. We found that ParB dimers are unable to engage with both halves of the palindromic *parS* site when adopting the predominant conformation with tethered N and M domains, ParB^Intra^ and ParB^Inter^. Partial unfolding of ParB appears to be needed for full engagement of *parS* DNA. This idea explains how *parS* DNA selectively stabilizes an intermediate of the reaction, while also binding to the reactant and product of the reaction, albeit with reduced affinity. Available co-crystal structures are consistent with the notion of a steric clash between *parS*-bound ParB chains. The structure of *H. pylori* ParB (lacking the C domain) bound to *parS* (PDB: 4UMK) (38) shows partially unfolded N domains (in four variations). Similarly, a C-terminally truncated variant of *C. crescentus* ParB (PDB: 6T1F) displays partial unfolding in one of the two *parS*-bound chains. These structures imply that *parS* DNA binding sufficiently compensates for the energetic investment in the partial unfolding of the N domain. The fact that a cysteine pair at the N-M interface (A102C/L134C) hinders sporulation and like another cysteine pair (A102C/ H133C) leads to elevated levels of the ParB^Inter^ conformation (Fig. S1C) supports the notion that this interface is indeed important for function and conformational control. We also demonstrated that the apposition of *parS* half sites is critical for catalysis. Spacer addition not only eliminated catalysis but lead to inhibition of the reaction, presumably due to improved binding of ParB^Intra^ dimers to the modified *parS* sites (Fig. 2B). This paradigm puts forward a strategy for engineering self-loading clamps with altered targeting specificity by repurposing sequence-specific or sequence-guided DNA binding proteins, eventually allowing for flexible chromosome labelling and DNA detection for diagnostic purposes.

### ParB CTP hydrolysis in partition complex assembly

We showed here that CTP hydrolysis by ParB is dispensable for some functions of ParABS in *B. subtilis* and critical for others. Sporulation and Smc recruitment were largely unperturbed in the CTP hydrolysis-deficient mutants E78Q and E111Q (Fig. 4). In contrast, ParB mutants interfering with CTP binding (G77S and R80A) are deficient in Smc recruitment (37, 50) as well as sporulation (28, 42). The robustness of Smc recruitment was surprising, particularly when considering the mediocre enrichment of ParB(E111Q) at *parS* sites. Conceivably, the poor enrichment is compensated by ParB(EQ) being trapped in a state that efficiently supports Smc recruitment. It is thus tempting to speculate that the CTP-ParB^Inter^ state is proficient in Smc loading and conceivably even solely responsible for it. In case of sporulation, any closed ParB clamps accumulating off the chromosome may also contribute to the conversion of ParA ATP-dimers to monomers—which is a prerequisite for sporulation—since a HTH mutant appears to support sporulation quite well (24). In contrast, chromosome segregation in the absence of *smc* was strongly compromised in either CTP-hydrolysis mutant, indicating defective ParABS function (Fig. 4). This defect likely explains why the CTPase activity is highly conserved and maintained over extended periods of time during evolution.

Explanations for the poor function of ParABS in the absence of CTP hydrolysis include (1) an inadequate assembly of the partition complex and (2) the accumulation of ParB in a state that is incompatible with proper ParABS function. We present evidence for either scenario. The notion of altered states of ParB in the EQ mutants is supported by our cysteine cross-linking (Fig. 6). While CTP hydrolysis-proficient wild-type ParB shows only low levels of ParB^Inter^ clamp formation *in vivo* (Fig. 1A, 6A), the EQ mutant variants predominantly adopt this state. It is also consistent with a recent report of CTP-controlled ParA-ATPase activation in the F plasmid ParABS system (51). Future work will have to establish, which form of ParB supports Smc loading and which one promotes ParA ATP hydrolysis. This may potentially establish temporal or spatial control of chromosome organization, segregation, and DNA replication.

Several observations point to defects in concentrating ParB in partition complexes in E78Q and E111Q. ChIP and fluorescence imaging revealed a reduced enrichment and broadened distribution of ParB(EQ) proteins at and near *parS* sites. CTP hydrolysis conceivably supports partition complex assembly by recycling ParB^Inter^ species. We envision two pools of ParB^Inter^ that may require recycling: ParB clamps that have excessively spread away from the *parS* loading site and clamps that have undergone N-gate closure without *parS* DNA stimulation, either becoming locked off the chromosome or trapped on the chromosome at large distances from *parS* sites (Fig. 7A). Such clamps would be permanently lost from the pool of productive ParB protein without the ability to hydrolyze CTP. *parS*-independent clamp closure readily occurs *in vitro* at least with *B. subtilis* ParB as indicated by the low but appreciable rates of CTP hydrolysis without *parS* DNA stimulation and by the slow but robust closure of the N-gate with the help of CTPγS in the absence of *parS* DNA. Similarly, we detected closed ParB(E78Q) clamps in cells lacking *parS* sites (Fig. 6B). Nevertheless, we cannot exclude the possibility that the poor enrichment at *parS* sites is caused at least in part by indirect consequences of the E78Q and E111Q mutations on ParB activity, for example by reducing N-gate stability or ParB autoinhibition.

We also described ParB behaviour by a simple reaction scheme (Fig. 7B). Thanks to CTP hydrolysis, ParB exhibits an enhanced affinity for *parS* sequences, beyond the one that would be possible at equilibrium (e.g. by hydrolysis deficient mutants). This ultra-affinity (52) corresponds to a roughly 50-fold reduction of the dissociation constant, from an estimate of the hydrolysis and *parS*-catalyzed ParB closure rates (*K*_*h*_ ∼1/ min; 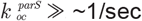; (37)). The enrichment of ParB on the *parS* flanking regions is a consequence of this effect because the increased amount of ParB bound to *parS* can subsequently diffuse away, thus populating the nearby non-specific DNA sequences.

Wild-type ParB barely adopts the clamp state *in vivo* despite being mostly localized in ParB foci (9, 53, 54). Mechanisms other than DNA entrapment may thus contribute to the retention of wild-type ParB protein in the partition complex, such as contacts between (open) ParB dimers (3, 28, 30, 55). DNA unloading upon CTP hydrolysis is prevented *in vitro* by the presence of CTP in case of *B. subtilis* ParB (Fig. 3F) as well as *C. crescentus* ParB (35). This implies that CDP is readily substituted for CTP without ParB unloading from the chromosome (Fig. 7B). It will be interesting to establish whether this nucleotide exchange reaction requires localization of ParB to the partition complex or DNA entrapment by ParB. If so, this feature would selectively eliminate off-target clamps or DNA-free clamps, respectively, thus further enhancing *parS*-proximal enrichment of ParB.

We conclude that *parS* DNA catalysis and ParB CTP hydrolysis lead to ParB/*parS* ultra-affinity and super-concentrate ParB within the partition complex. The unusually high concentration of ParB (56) is likely required to optimally support chromosome and plasmid segregation by the ParABS diffusion-ratchet mechanism but might not be essential for other cellular functions of ParABS such as Smc recruitment.

## Materials and Methods

### *Bacillus subtilis* strains and growth

The *B. subtilis* 1A700 isolate was used for all experiments. *B. subtilis* was transformed via natural competence and homologous recombination as described in (57) and grown on SMG-agar plates with appropriate antibiotic selection. Transformants were next checked by PCR and Sanger sequencing. Tagging of ParB (with HaloTag or mScarlet) or HBsu (with mTurquoise) proteins was done at the carboxy terminus. Genotypes of strains used in this study are listed in Table S1.

For spotting assays, the cells were cultured in SMG medium at 37 °C to stationary phase and 9^2^ and 9^5^-fold dilutions were spotted onto ONA (∼16 h incubation) and SMG (∼24 h incubation) agar plates.

### Expression and purification full-length *B. subtilis* ParB protein

Expression constructs were prepared in pET-28 derived plasmids by Golden-Gate cloning. Untagged recombinant proteins were produced in *E. coli BL21-Gold (DE3)* grown in ZYM-5052 autoinduction media at 24 °C for 24 hours. Purification of full-length ParB protein was done as described before in (soh et al., 2019). In brief, cells were lysed by sonication in buffer A (1 mM EDTA pH 8, 500 mM NaCl, 50 mM Tris-HCl pH 7.5, 5 mM β-mercaptoethanol, 5 % (v/v) glycerol, and protease inhibitor cocktail (PIC, Sigma)). Ammonium sulfate was added to the supernatant to 40 % (w/v) saturation and kept at 4 °C for 30 minutes while stirring. The sample was then centrifuged, the supernatant was collected, and ammonium sulfate was added to 50 % (w/v) saturation and kept at 4 °C for 30 minutes while stirring. The pellet was then collected by centrifugation and dissolved in buffer B (50 mM Tris-HCl pH 7.5, 1 mM EDTA pH 8 and 2 mM β-mercaptoethanol). The sample was additionally diluted with buffer B to a conductivity of 18 mS/cm and loaded onto a Heparin column (GE healthcare). The protein was eluted with a linear gradient of buffer B containing 1 M NaCl. Peak fractions were collected and diluted with buffer B to a conductivity of 18 mS/cm and loaded onto HiTrap SP columns (GE healthcare). A linear gradient of buffer B containing 1 M NaCl was used for elution. Peak fractions were collected. and directly loaded onto a Superdex 200 16/600 pg column (GE healthcare) preequilibrated in 300 mM NaCl and 50 Mm Tris-HCl pH 7.5. For cysteine mutant ParB, 1 mM TCEP was added to the gel-filtration buffer. For ITC measurements, peak fractions were collected, diluted 1:1 with buffer containing 50 mM Tris-HCl pH 7.5 and 10 mM MgCl2 to bring the final buffer to 150 mM NaCl, 50 mM Tris-HCl pH 7.5 and 5 mM MgCl2 and directly used for measurements.

### CTPγS

CTPγS was custom-synthesized and purified by reversed-phase chromatography (RP-HPFL) to a final purity of 90.6 % by Jena Biosciences (Jena, Germany). A stock solution at a concentration of 100 mM (pH-adjusted to 8.0 by addition of NaOH) was aliquoted and stored at -80 °C.

### Isothermal titration calorimetry (ITC)

The measurement was done using MicroCal iTC200 (GE Healthcare Life Sciences). The instrument was pre-cooled to 4 °C. All measurements were made in buffer containing 150 mM NaCl, 50 mM Tris/HCl pH 7.5, and 5 mM MgCl2. Both measurement cell and injection syringe were subjected to a series of washes with buffer. 280 μL of the protein solution at 80-120 μM monomer concentration was added to the measurement cell, and the injection syringe was filled with buffer containing 2 mM of CTP or buffer only. Measurements were taken with an initial delay of 180 seconds, and the settings of the instrument were adjusted to: reference power of 5 μcal/ sec, a stirring velocity of 1000 RPM, and a “high feedback” mode. Raw data were integrated to kcal/mol, presented as a Wiseman plot, and wherever possible, regression curves were calculated according to a 1:1 nucleotide to ParB monomer binding model. Origin (GE Healthcare) was used for fitting results from the measurements by using the following equation:

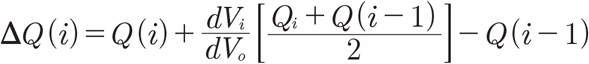

*V*_*i*_ is the injection volume of ligand (nucleotides), *V*_*o*_ is the volume of the cell, *Q(i)* is the heat released from the i^th^ injection which is in turn calculated using the following equation:

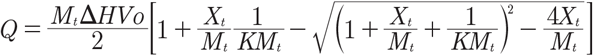

*K* is the binding constant, *ΔH* is the molar heat of ligand binding, *Xt* is bulk concentration of nucleotide, and *Mt* is the bulk concentration of ParB (moles/liter) in *Vo. K* and *ΔH* were estimated by Origin, and *ΔQ(i)* for each injection was calculated and compared to the measured heat. *K* and *ΔH* estimates were then improved using standard Marquardt methods. Several iterations were performed until the fit could no longer be improved.

### Measurement of CTP hydrolysis by Malachite Green colorimetric detection

Mixtures of CTP (2x) with or without ^*parS*^DNA _40_ (2x) in reaction buffer (150 mM NaCl, 50 mM Tris pH 7.5, 5 mM MgCl2) were prepared on ice. Protein solutions (2x) in reaction buffer were also prepared on ice. CTP/^*parS*^DNA _40_ pre-mix (10 μL) was added to protein solution (10 μL) using BenchSmart 96 (Rainin) dispenser robot and mixed by pipetting. After mixing, samples (containing CTP 1 mM, ^*parS*^DNA _40_ 1 μM, protein 10 μM) were placed in a PCR machine and set to incubate at 25 °C for 1 hour. In parallel, phosphate blanks were prepared for each experiment. Samples were diluted 4-fold by the addition of 60 μL water, then mixed with 20 μL working reagent (Sigma) and transferred to a flat bottom 96 well plate. The plate was left to incubate for 30 minutes at 25 °C, and the absorbance was then read at a wavelength of 620 nm. Absorbance values from the phosphate standard samples were used to plot an OD_620_ versus phosphate concentration standard curve. Raw values were converted to rate values using the standard curve. Absolute rates were calculated by normalizing for protein concentration. Mean values and standard deviation were calculated from three repeat experiments. Graphs were plotted on GraphPad Prism for presentation.

### 40-nucleotide, double stranded ^*parS*^DNA _40_ preparation

Complementary strands of oligonucleotides (at 100 μM each) (for sequences see Supplementary Table S2) were mixed 1:1 and heated to 95 °C for 10 minutes and then left to cool down to 25 °C.

### *In vitro* cross-linking

Mixtures (2x) of NTP with or without ^*parS*^DNA_40_ were prepared in reaction buffer (150 mM NaCl, 50 mM Tris-HCl pH 7.5, 5 mM MgCl2) and kept at room temperature (RT) for 5 minutes. Protein solution (2x in 300mM NaCl, 50mM Tris-HCl pH 7.5) was added to a final concentration of 10 μM and the mixture (containing 1 mM NTP with or without 1 μM ^*parS*^DNA_40_ was incubated for an additional 5 minutes at room temperature. 1 mM BMOE (from a 20 mM stock solution) was added to the samples. After 5 minutes at RT, samples were quenched with β-mercaptoethanol (23 mM final), loading dye was added, and samples were incubated at 70°C for 5 minutes and loaded onto Bis-Tris 4-12 % gradient gels (ThermoFisher). Bands were stained by CBB and relative band intensity was quantified by scanning and semi-automated analysis in ImageQuant (GE Healthcare).

### *In vivo* cross-linking

Culture, grown overnight in 0.5 % (w/v) glucose containing LB, was used to inoculate fresh LB medium to an OD_600_ of 0.005 and were grown at 37°C to mid-exponential phase (OD_600_of 0.3). Around 70g of Ice cubes were added directly to the culture flask, and the cells were harvested by centrifugation for 5 minutes at 14,000 g at 4°C. The cell pellet was washed in cold PBSG (PBS with 0.1 % (v/v) glycerol). Cells were pelleted again and resuspended in 1 mL cold PBSG. 1.25 OD units were collected, and resuspended in 30 μL of cold PBSG. 0.5 mM BMOE was then added to the samples, mixed by pipetting, and placed on ice for 15 minutes. 15 mM β-mercaptoethanol was added and the samples were incubated on ice for 3 minutes to quench the reaction. The following reagents were added to the given final concentrations: 750 U/mL Benzonase (Sigma), 5 μM HaloTag-TMR (Promega), 47 U/μL Ready Lyse Lysozyme (Epicentre), and 1x PIC (Sigma). The cells were then lysed at 37°C in the dark for 30 minutes, loading dye containing 5 % (v/v) β-mercaptoethanol was added, and the samples were incubated at 70 °C for 5 minutes. 10 μL of each sample was loaded onto a 4-12 % Bis-Tris Gel (ThermoFisher). Following SDS PAGE, gels were imaged on an Amersham Typhoon (GE Healthcare) with Cy3 DIGE filter setup. Bands were quantified using ImageQuant (GE Healthcare).

### Chromosome Entrapment Assay

The assay was done as described in (48, 49) using agarose beads. In brief, cells were grown, harvested, and 3.75 OD units was cross-linked as described above (*in vivo* cross-linking). The sample was split into two aliquots (2:3 for the output/beads and 1:3 for input). To each input sample, 5 µL of Mastermix-1 was added. Mastermix-1 contains (per one input sample): 400 units of Ready-Lyse Lysozyme, 12.5 units Benzonase, 1 µM TMR HaloTag, 0.5 µL protease inhibitor cocktail (PIC), and 0.9 µL 1X PBS). The input samples are then incubated for 25 min at 37 °C covered from light, and an equal volume of 2X protein loading dye was added and the samples were stored at -20 °C.

For agarose bead preparation, 1 µL PIC and 9 µL of Dynabeads™ M-280 Streptavidin (ThermoFisher Scientific) were added to each output (‘beads’) sample. 100 µL of preheated 2 % low melt agarose was then added, and immediately followed by 700 µL of mineral oil (preheated to 4 °C). The sample was then vortexed vigorously for 2 min on ice. Each sample was treated quickly and separately.

The output (beads) samples were then spun down at 10,000 g at RT, and the supernatant (oil) was removed as much as possible. The beads were then resuspended in 1 mL PBSG (0.1 % glycerol) very gently to prevent the release of cells from the agarose beads.

To each output sample, 4U/µL of Ready-Lyse Lysozyme, 1 µM of TMR-HaloTag, 5 µL PIC, and 1 mM EDTA were added (same as Mastermix-1 but without Benzonase). The output samples are then incubated for 25 min at 37 °C covered from light. Next, the beads were washed twice with 1 mL RT PBSG, and with TES buffer (50 mM Tris-HCL pH 7.5, 10 mM EDTA, 1 % SDS) three times, with the first time incubated for 1 hr at RT with moderate shaking (500 rpm) and the following two incubations were done for 30 minutes. The beads were resuspended in 1 mL TES and incubated overnight at 4 °C on a rotating wheel and covered from light.

The next day, the beads were washed twice with PBS 1X at RT to remove EDTA and SDS and resuspended in 100 µL of PBS. 1µL of SmDnase (750 U) was added to each output sample and incubated for 1 hr at 37 °C with moderate shaking and covered from light. The samples were next incubated for 1 min at 70 °C at 14000 rpm shaking to melt the agarose plugs, and then transferred onto ice and incubated for 5 min. The samples were spun down at maximum speed for 15 min at 4 °C and the supernatant was transferred to an acetate spin column and spun down for additional 5 min at RT at maximum speed to recover as much liquid as possible. The approximate recovered volume is 100 µL which was then diluted to 1 mL with water and 10 µL of 2 % sodium-deoxycholate and 3.3 mg/mL BSA was added and incubated for 30 minutes at 4 °C protected from light. After incubation, 120 µL of 80 % trichloroacetic acid (TCA) solution was added and incubated for 2 hours on ice and covered from light. The samples are then spun down for 15 minutes at maximum speed at 4 °C. The precipitate was collected and resuspended in 10 µL of 1x loading dye. The input samples were thawed and boiled with the output samples for 5 min at 95 °C. 5 % of the input and all of the output volume were loaded on an SDS-PAGE using 8-12 % Novex Wedge Well Tris-Glycine gel (Thermo Scientific). The gel was then imaged on Amersham Typhoon (GE Healthcare) with Cy3 filter.

### Biolayer interferometry (BLI)

Measurements were done in a buffer containing 150 mM NaCl, 50 mM Tris-HCl pH 7.5, and 5 mM MgCl2 on BLItz machine (FortéBio Sartorius). Streptavidin coated biosensors were used in all the measurements and were hydrated in the reaction buffer for 10 minutes before loading. A baseline was first recorded by equilibrating the biosensor in 250 µL reaction buffer in a black 0.5 mL Eppendorf tube for 30 seconds. 4 µL of 100 nM biotin labelled double stranded *parS* or *mut-parS* DNA^169bp^ was then loaded on the biosensor for 5 minutes. After the DNA loading phase, the biosensor was washed once with the reaction buffer and once with the reaction buffer containing 1 mM NTP. Next, 2X ParB solution and 2x NTP solution were mixed 1:1 (final concentration of 1 µM parB and 1 mM NTP) and 4 µl of the mixture was loaded immediately on the biosensor for 2 minutes. The dissociation phase is then carried for 5 minutes in 250 µL protein-free reaction buffer with or without NTP. All measurements were analyzed on the BLItz analysis software and replotted on GraphPad Prism for presentation.

### Chromatin Immunoprecipitation (ChIP)

ChIP samples were prepared as described previously (Bürmann et al., 2017) with minor modifications. Cells were grown in 200 mL minimal media (SMG) at 37°C until mid-exponential phase (OD600=0.03). Next, 20 mL of fixation buffer F was added (50 mM Tris-HCl pH 7.4, 100 mM NaCl, 0.5 mM EGTA pH 8.0, 1 mM EDTA pH 8.0, 10% (w/v) formaldehyde) for 30 minutes at RT with occasional shaking. Cells were harvested by filtration and washed in 1X PBS. Each sample was adjusted for 2 OD600 units (2 mL at OD600 = 1) and resuspended in TSEMS lysis buffer (50 mM Tris pH 7.4, 50 mM NaCl, 10 mM EDTA pH 8.0, 0.5 M sucrose and PIC (Sigma), 6 mg/mL lysozyme from chicken egg white (Sigma)). After 30 minutes of incubation at 37 °C with vigorous shaking protoplasts were washed again in 2 mL TSEMS, resuspended in 1 mL TSEMS, split into 3 aliquots, pelleted, flash frozen, and stored at -80°C for further use.

For ChIP-qPCR, each pellet was resuspended in 2 mL of buffer L (50 mM HEPES-KOH pH 7.5, 140 mM NaCl, 1 mM EDTA pH 8.0, 1 % (v/v) Triton X-100, 0.1 % (w/v) Sodium-deoxycholate, 0.1 mg/mL RNaseA and PIC (Sigma)) and transferred to 5 mL round-bottom tubes. Cells were sonicated three times for 20 sec on a Bandelin Sonoplus with a MS72 tip (settings: 90 % pulse and 35 % power output). The lysates were next transferred into 2 mL tubes and centrifuged for 10 min at 21000 g at 4°C. 800 µL of the supernatant was used as input (IP) and 200 μL was kept as whole cell extract (WCE). The WCE tubes are frozen at -20°C for later.

For the IP, antibody serum was pre-incubated with Protein G coupled dynabeads (Invitrogen) in 1:1 ratio for 2h at 4°C on a rotating wheel. Next, the beads were washed in buffer L and 50 μL were aliquoted to each sample tube. Samples were incubated with the beads for 2h at 4°C with rotation. After incubation, all samples were subjected to a series of washes with buffer L, buffer L5 (buffer L containing 500 mM NaCl), buffer W (10 mM Tris-HCl pH 8.0, 250 mM LiCl, 0.5 % (v/v) NP-40, 0.5 % (w/v) Na-Deoxycholate, 1 mM EDTA pH 8.0) and buffer TE (10 mM Tris-HCl pH 8.0, 1 mM EDTA pH 8.0). After washing, the beads were resuspended in 520 µL buffer TES (50 mM Tris-HCl pH 8.0, 10 mM EDTA pH 8.0, 1 % (w/v) SDS). The WCE are thawed and 300 µL of TES and 20 µL of 10 % SDS were also added. Both tubes were incubated O/N at 65°C with vigorous shaking to reverse the formaldehyde cross-linking. The next day, Phenol-chloroform DNA extraction was performed to purify the de-cross-linked DNA. Samples were transferred to screw cap 1.5 mL tubes and mixed vigorously with 500 μL of phenol equilibrated with buffer (10 mM Tris-HCl pH 8.0, 1 mM EDTA). After centrifugation (10 minutes, RT, 13000rpm), 450 µL of the aqueous phase was transferred to a new screw cap tube and mixed with equal volume of chloroform, followed by centrifugation. 400 µL of aqueous phase was then recovered for DNA precipitation with 1 mL of 100% ethanol (2.5x volume), 40 µL of 3M NaOAc (0.1x volume), and 1.2 µL of GlycoBlue and incubated for 20 minutes at -20°C. Lastly, samples were centrifuged for 10 minutes at 20000g at RT and the pellets were purified with a PCR purification kit, eluting in 50 μL buffer EB.

For qPCR, 1:10 and 1:1000 dilutions in water of IP and WCE were prepared, respectively. Each 10 µL reaction was prepared in duplicate (5 µL Takyon SYBR MasterMix, 1 µL 3 µM primer pair, 4 µL of DNA) and run in Rotor-Gene Q machine (QIAGEN). Primer sequences are listed in the table S2. Raw Data were analyzed using PCR Miner server (http://ewindup.info) (Zhao & Fernald, 2005).

For IP of samples for ChIP-Seq, the procedure was the same as for ChIP-qPCR, except for resuspending the pellets in 1 mL of buffer L and sonication in a Covaris E220 water bath sonicator for 5 min at 4°C, 100 W, 200 cycles, 10% load and water level 5.

For deep sequencing, the DNA libraries were prepared by Genomic Facility at CIG, UNIL, Lausanne. Briefly, the DNA was fragmented by sonication (Covaris S2) to fragment sizes ranging from 220–250 bp. DNA libraries were prepared using the Ovation Ultralow Library Systems V2 Kit (NuGEN) including 15 cycles of PCR amplification. 80 to 100 million single end sequence reads were obtained on a HiSeq4000 (Illumina) with 151 bp read length.

### Processing of ChIP-Seq reads

Reads were mapped to *Bacillus subtilis* genome NC_000964.3 with bowtie2 using the default mode. Subsequent data analysis was performed using Seqmonk http://www.bioinformatics. babraham.ac.uk/projects/seqmonk/. A bin size of 1 kb was used.

### Mathematical model of ParB-DNA association

The steady state solution of the model in Fig. 7B relies on the absence of net fluxes over the reaction network. Thus, the concentration of unbound ParB in the open state is related to the concentration of unbound ParB in the closed state by the relation

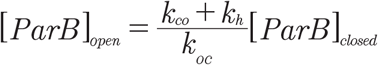

The total amount of unbound ParB is

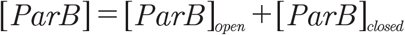

which can be used to write

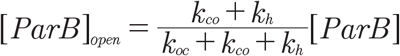

Similarly,

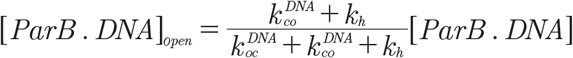

where [ParB·DNA] is the total concentration of ParB bound to DNA. Here we assumed that the hydrolysis rate is not significantly affected by the bound DNA.

From these relations it is possible to determine the observed dissociation constant of ParB (independently of its open or closed state) for DNA:

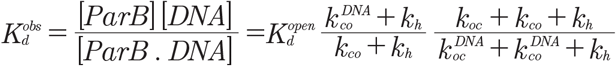

There are three cases:

1) *k*_*h*_*≈ 0*: Hydrolysis deficient mutant

The dissociation constant reduces to

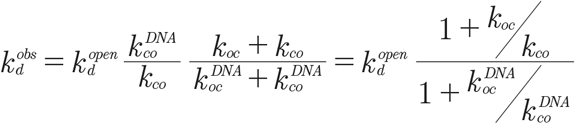

If DNA (whether *parS* of non-specific sequences) only accelerates the transitions between the open and closed states but does not alter significantly their equilibrium, then

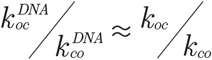

and consequently

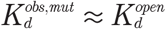

2) non-specific DNA

In this case the transitions between the open and closed states is unaffected by substrate binding:

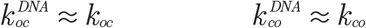

and again, as a consequence

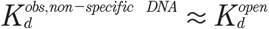

3) *parS* DNA:

We know that

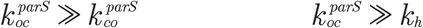

the first relation is due to the spontaneous closure of hydrolysis-deficient mutants (with CTP) or wild-type protein (with a non-hydrolysable analogue) (Fig. S4), telling that the closed state is lower in free energy than the open state; the second obtained from the observation that ∼40-fold sub-stoichiometric amounts of *parS* DNA are sufficient to fully stimulate CTP hydrolysis and N-gate closure, meaning that a step other than closure (presumably hydrolysis) is rate-limiting (37). Furthermore, considering that for free ParB it is *k*_*h*_ *>> k*_*co*_,*k*_*oc*_ to ensure proper recycling, we have

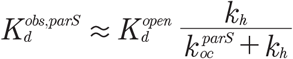

which is much smaller than 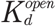.

## Acknowledgements

We are grateful to members of the Gruber lab for stimulating discussions and comments on the manuscript and to Tung Le for sharing unpublished results.

## Supplementary Materials for

**Figure S1.**
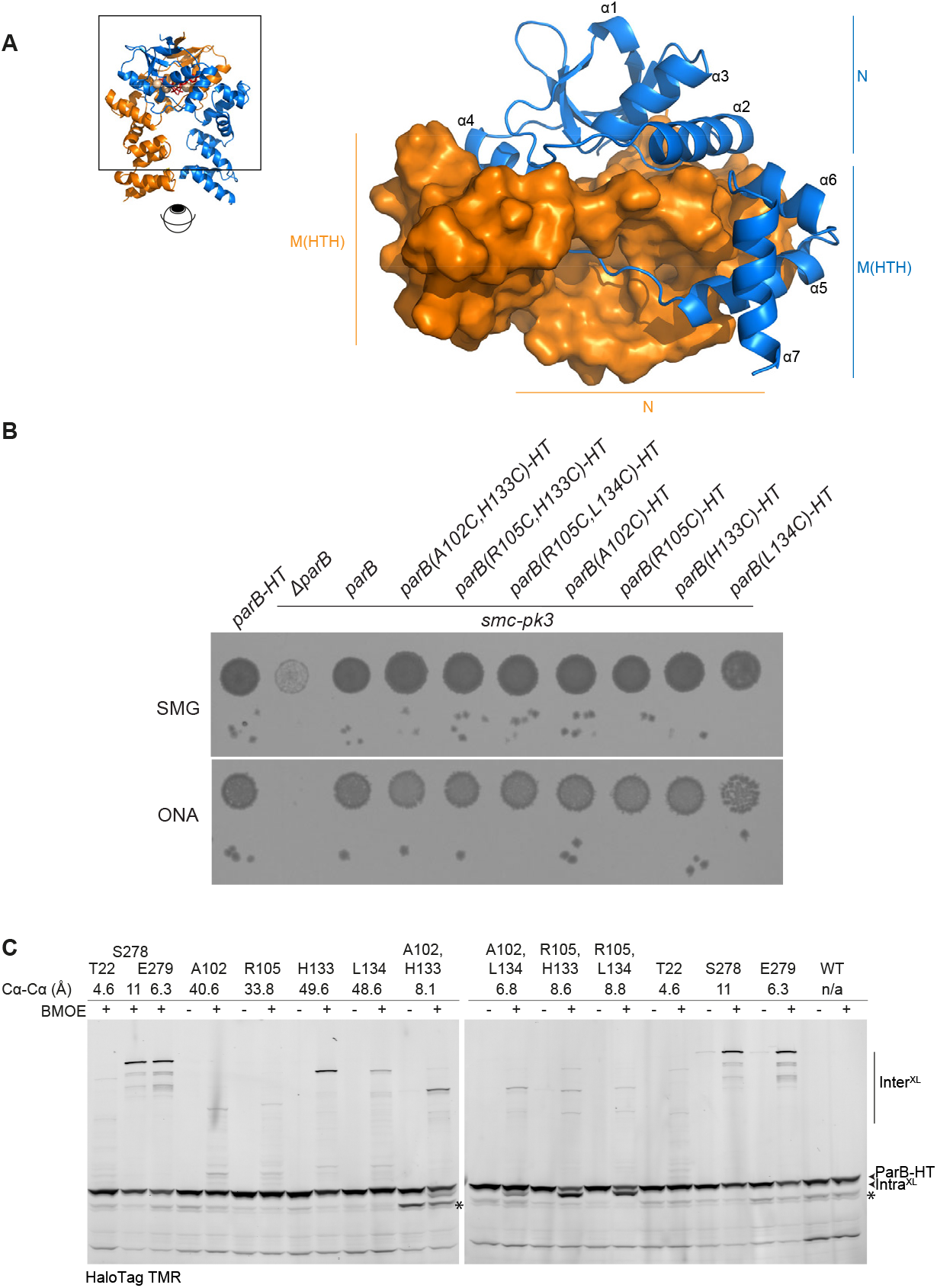
**(A)** ParB-CDP crystal structure (PDB: 6SDK) showing the tight interlocking between N and M domains from the two chains of a ParB dimer as viewed from the bottom. **(B)** Growth assay by dilution spotting of strains harbouring different *parB* alleles combined with *smc-pk3*. 9^2^ and 9^5^-fold dilutions were spotted on minimal medium agar plates (SMG) and rich medium agar plates (ONA) and imaged after 24 and 16 hours, respectively. **(C)** *In vivo* BMOE cross-linking of selected ParB cysteine mutants. Same as in Fig. 1A with additional strains and control reactions lacking BMOE. For reference, the Cα-Cα distances for pairs of residues on the ParB-CDP dimer structure (PDB: 6SDK) are given. Asterisks denote degradation products of ParB-HT.

**Figure S2.**
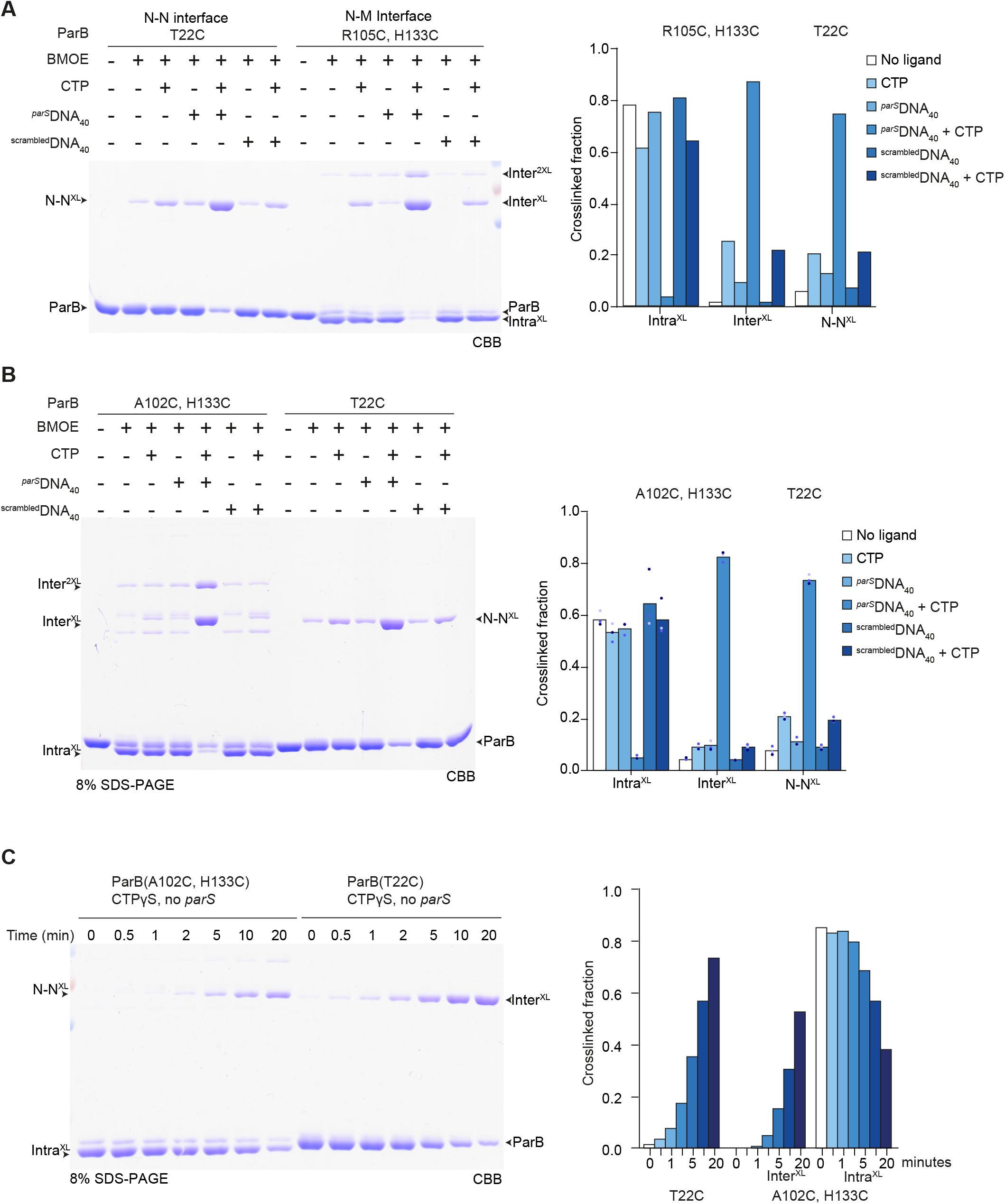
**(A)** *In vitro* BMOE cross-linking of purified cysteine mutants ParB(T22C) and ParB(R105C, H133C). Cross-linked fractions were analysed by SDS-PAGE and CBB staining. Same gel picture as shown in Fig. 1B with estimation of cross-linking efficiencies from quantification of gel band intensities (right panel). **(B)** *In vitro* BMOE cross-linking of purified mutants as in (A) but using ParB(T22C) and ParB(A102C, H133C). Quantification of cross-linking efficiency was done in three replicates. Mean and individual data points are shown. **(C)** Time course of *in vitro* BMOE cross-linking of purified cysteine mutants ParB(A102C, H133C) and ParB(T22C) after addition of 1 mM CTPγS (in the absence of *parS* DNA). Quantification as in (A) (right panel).

**Figure S3.**
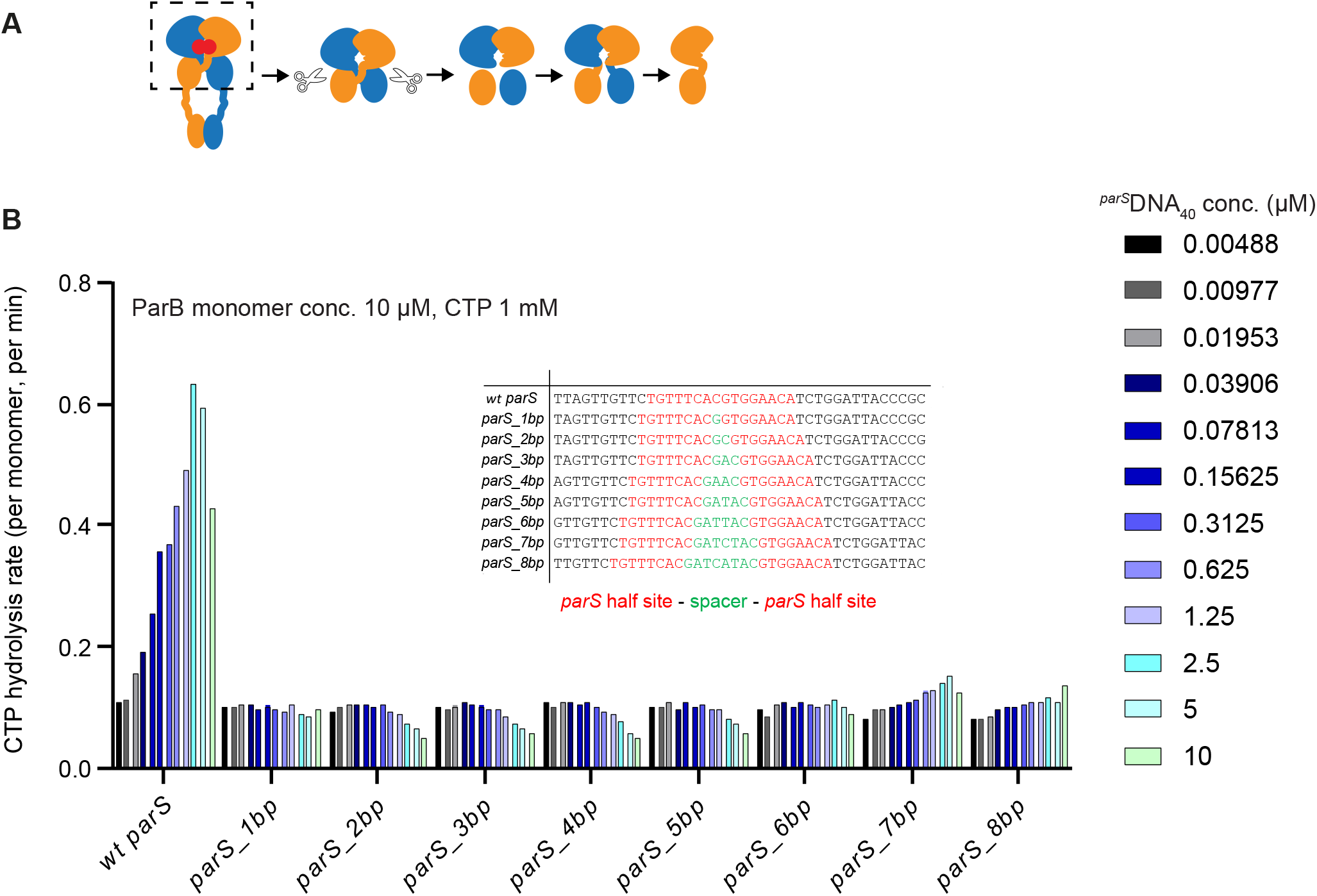
**(A)** A model of the ParB^Intra^ monomer was created manually by linking residues 1-116 from chain A (PDB: 6SDK) to residues 117-218 of chain B. **(B)** Rate of CTP hydrolysis by wild-type *B. subtilis* ParB measured by Malachite green assay. The rate was measured in response to the presence of different ^*parS*^DNA_40_ sequences with increasing number of spacer base pairs between the two *parS* half sites. Final reaction contained 10 µM of ParB monomer, 1 mM CTP, and a serial dilution of altered *parS* DNA, in Mg2+ containing buffer.

**Figure S4.**
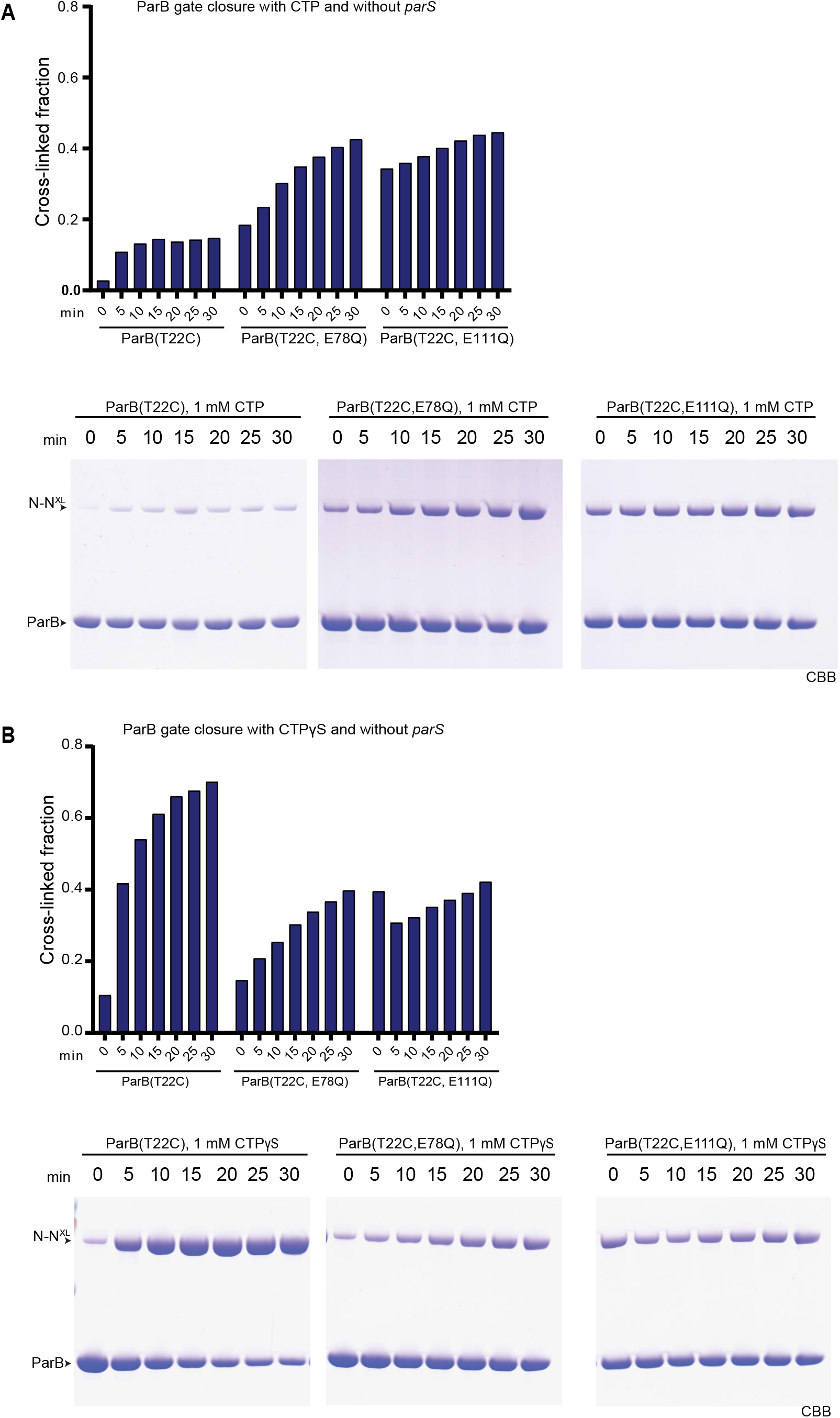
**(A)** Time course of *in vitro* BMOE cross-linking of ParB variants (10 µM), in the presence of CTP (1 mM) without *parS* DNA. Reaction is in Mg2+ containing buffer (see Materials and Methods). Quantification of bands was performed with ImageQuant (GE Healthcare). **(B)** Same as in (A) but with CTPγS (1 mM).

**Figure S5.**
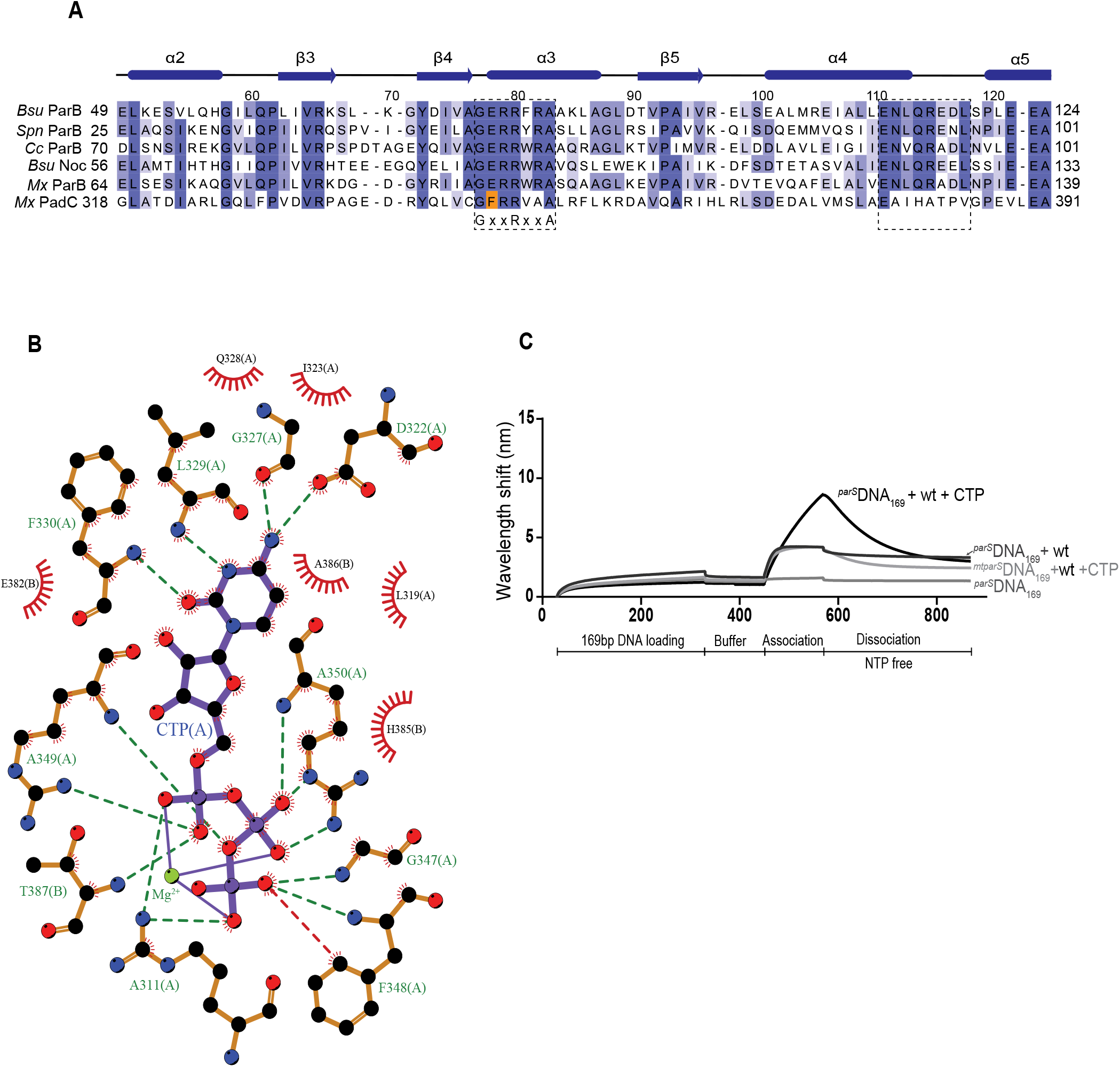
**(A)** Alignment of central sequences of four bacterial ParB proteins (*Bsu, Bacillus subtilis*; *Spn, Streptococcus pneumoniae; Cc, Caulobacter crescentus; Mx, Myxococcus xanthus*), *Bsu* Noc and *Mx* PadC. The conserved GE_78_RRxxA and E_111_NLQRE motifs are marked by dashed boxes. The change of nucleotide (E78 to F348) in *Mx* PadC is marked in orange colour. **(B)** 2D Protein-Ligand interaction map of PadC-CTP (PDB: 6RYK) showing residues from the two chains of PadC that contribute to the binding of one CTP molecule and one Mg2+. Hydrogen bonds are represented by green dashed lines and hydrophobic interactions by red semi-circles (except for F348(A) extra red dashed line is added). The map was generated on LigPlot+ software (58). **(C)**Biolayer interferometry assay measuring ParB loading efficiency on *parS* or *mtparS* 169 bp DNA in the presence or absence of CTP in the protein association phase. A negative control with no protein is also included (^*parS*^DNA_169_).

**Figure S6.**
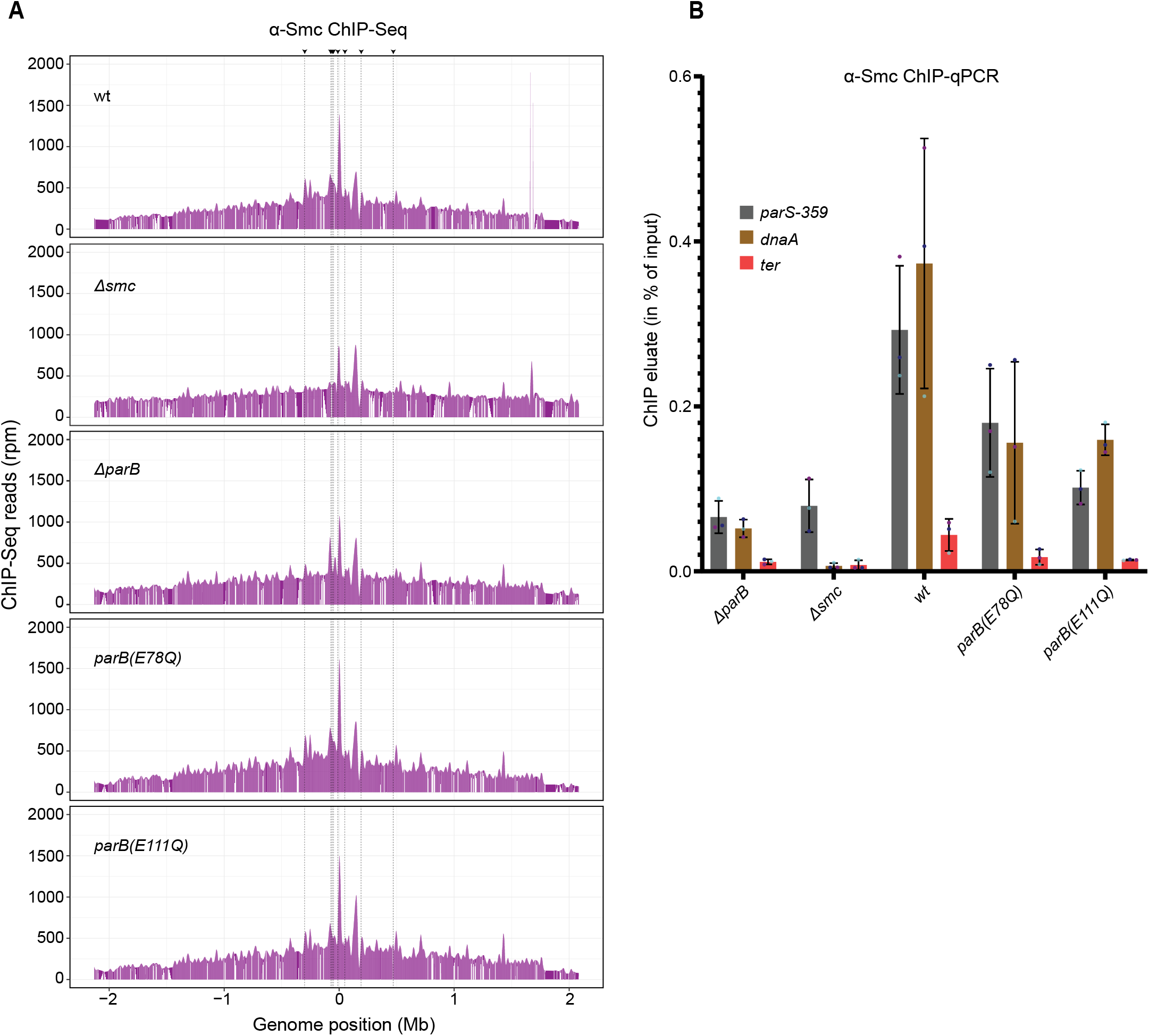
**(A)** Chromatin-immunoprecipitation coupled to deep sequencing (ChIP-Seq) using α-Smc serum. Panels show genome wide distribution of Smc enrichment. Same conditions in Fig. 3C but with the addition of *Δsmc* strain. **(B)** Chromatin-immunoprecipitation coupled to quantitative PCR (ChIP-qPCR) using α-Smc serum. Mean values and standard deviation from three repeat measurements are reported. Individual data points are shown as dots. Enrichment of Smc was tested at the three loci: *parS-359, dnaA*, and the replication terminus (*yocGH*).

**Figure S7.**
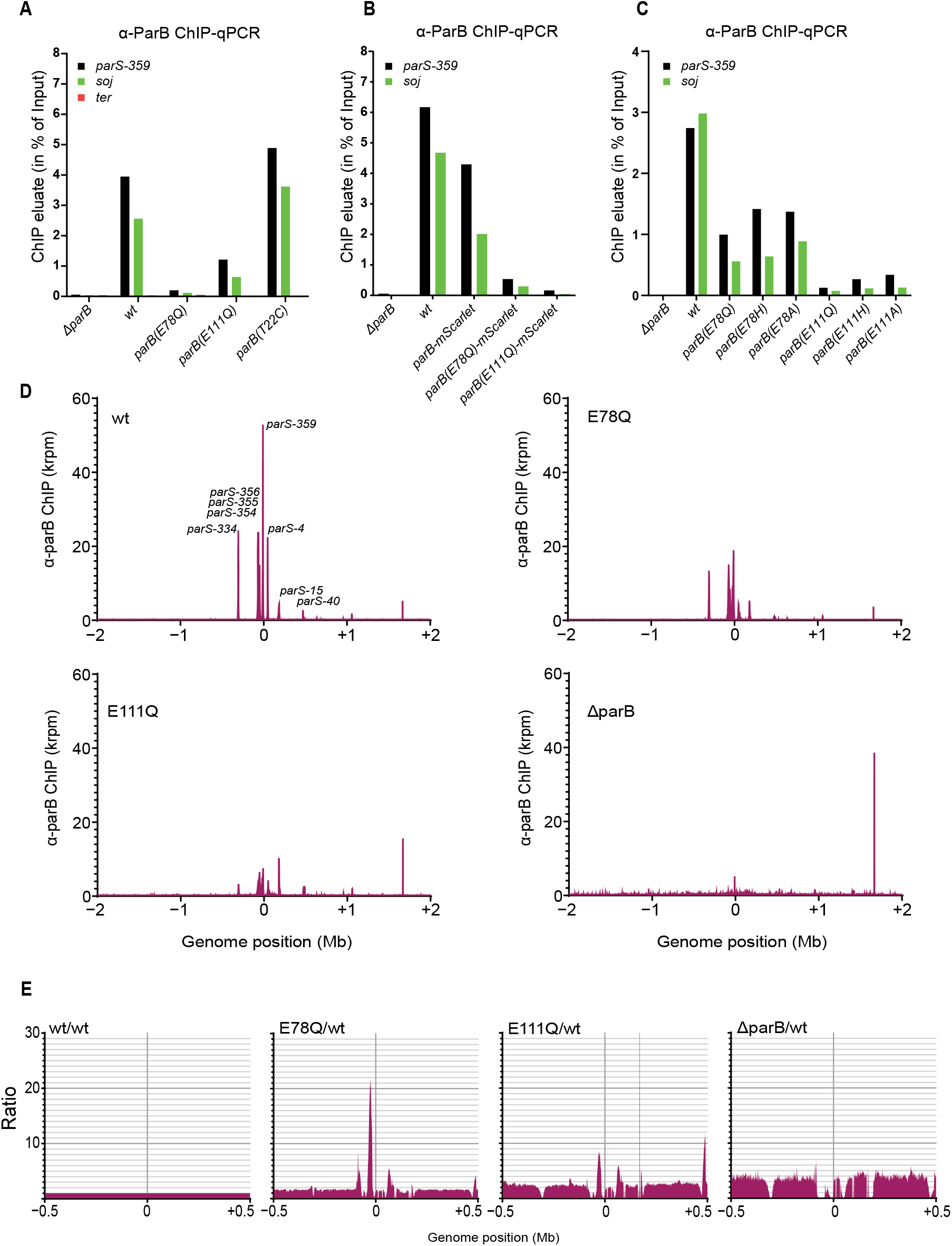
**(A)** Chromatin-immunoprecipitation coupled to quantitative PCR (ChIP-qPCR) using α-ParB serum. Same as in Fig. 5A in addition to a strain carrying the *parB(T22C)* allele. **(B)** Chromatin-immunoprecipitation coupled to quantitative PCR (ChIP-qPCR) using α-ParB serum for strains with mScarlet tagged ParB. As in (A). **(C)** Chromatin-immunoprecipitation coupled to quantitative PCR (ChIP-qPCR) using α-ParB serum. Same as in Fig. 5A in addition to strains carrying histidine (H) and alanine (A) substitutions of E78 and E111 residues. **(D)** Chromatin-immunoprecipitation coupled with deep sequencing (ChIP-Seq) using α-ParB serum. Same as in Fig. 5B but with the addition of a *ΔparB* strain and showing genome wide distribution. Of note, the peak at position +1.8 Mb represents enrichment at the *smc* gene presumed to be a contamination. **(E)** Ratio plot of sequencing reads found in E78Q, E111Q, and *ΔparB* divided by read number in wild type. The region shown is 1 Mb wide surrounding the origin of replication. Peaks correspond to regions with higher enrichment of ParB(EQ) near *parS* sites (higher spreading efficiency).

**Figure S8.**
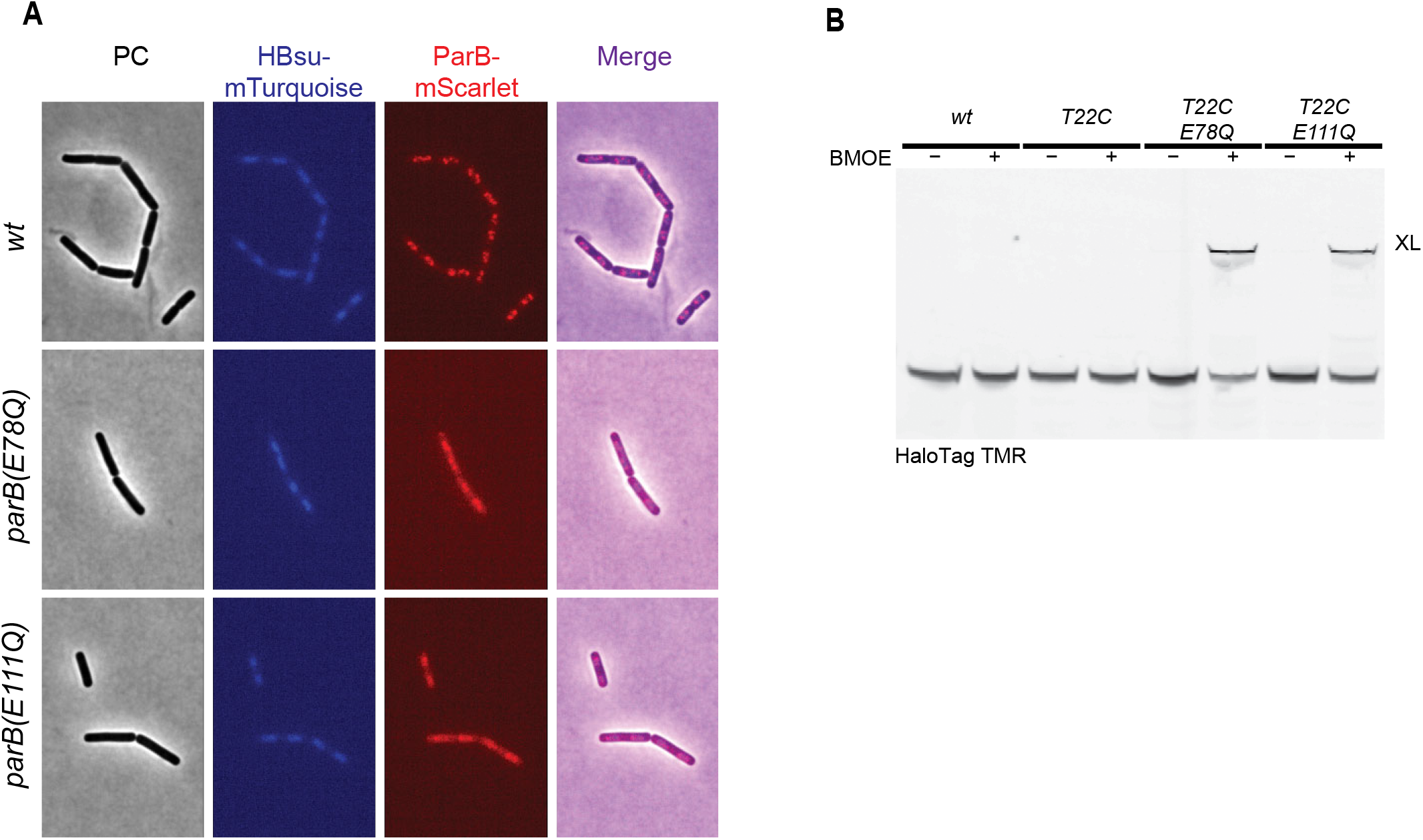
**(A)** Fluorescence microscopy imaging of *B. subtilis:* mScarlet tagged ParB (red) and mTurquoise tagged HBsu (blue). Same as in Fig. 5C but cells were pre-incubated with 200 µg/ml of chloramphenicol for artificial nucleoid compaction for 15 minutes prior to imaging. The images show colocalization of ParB-mScarlet signal with nucleoids staining. **(B)** Expression levels of ParB-HT variants as judged by HaloTag-TMR labelling and in-gel fluorescence detection. ParB-HT variants harbor the T22C residues and the E78Q or E111Q mutation.

**Table S1.**
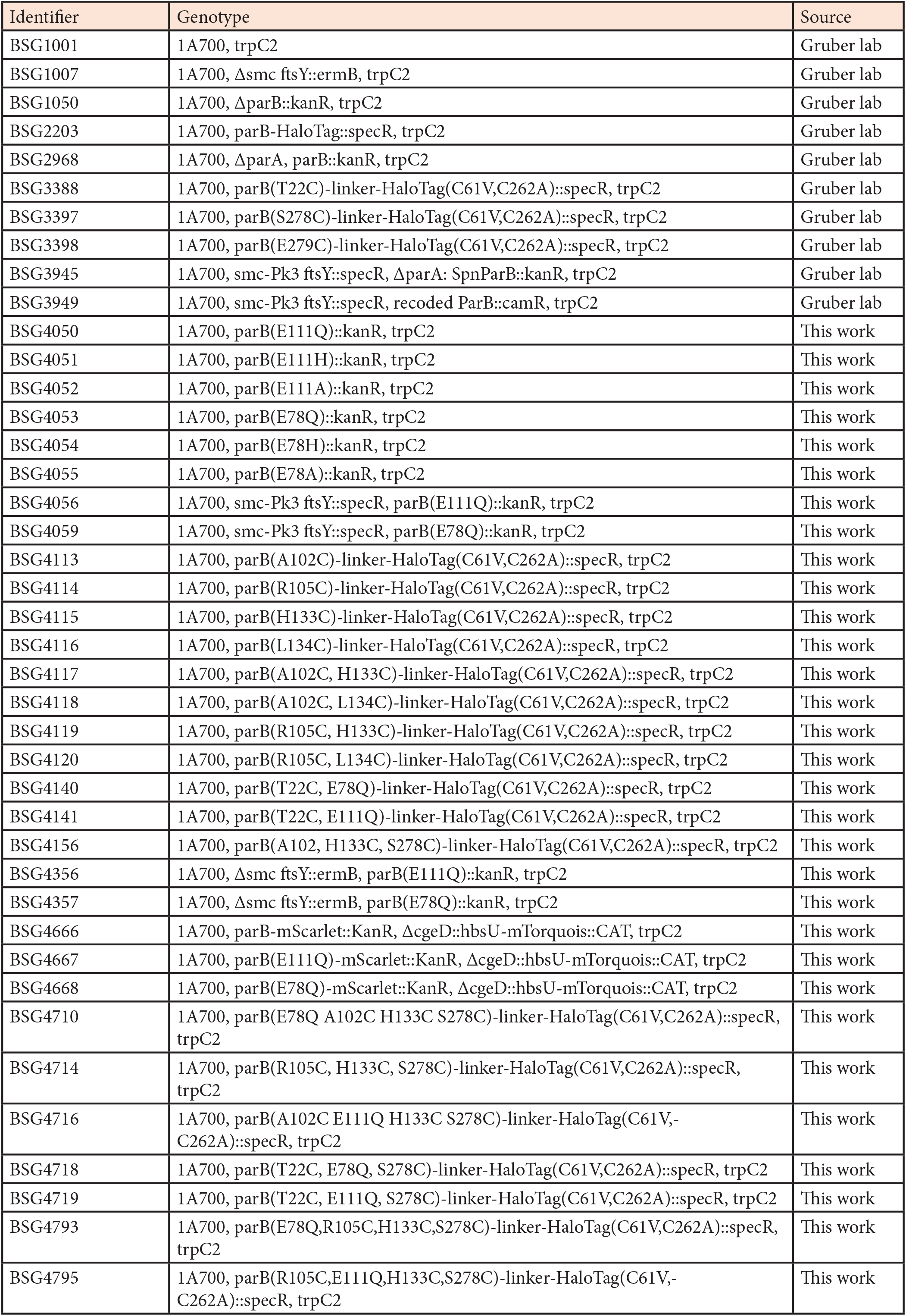
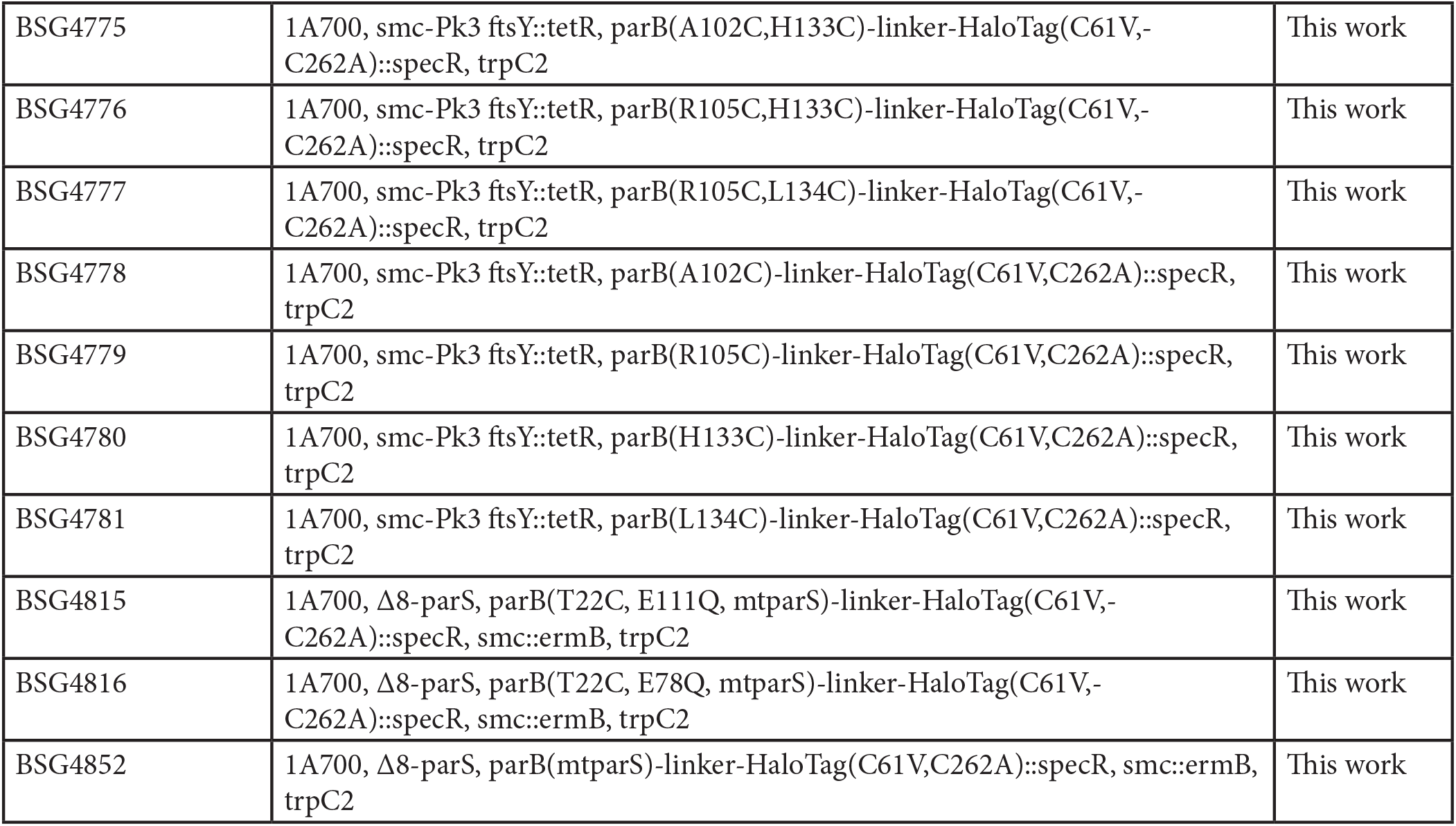
Strains used in this study.

| Identifier | Genotype | Source |
| --- | --- | --- |
| BSG1001 | 1A700, trpC2 | Gruber lab |
| BSG1007 | 1A700, Δsmc ftsY::ermB, trpC2 | Gruber lab |
| BSG1050 | 1A700, ΔparB::kanR, trpC2 | Gruber lab |
| BSG2203 | 1A700, parB-HaloTag::specR, trpC2 | Gruber lab |
| BSG2968 | 1A700, ΔparA, parB::kanR, trpC2 | Gruber lab |
| BSG3388 | 1A700, parB(T22C)-linker-HaloTag(C61V,C262A)::specR, trpC2 | Gruber lab |
| BSG3397 | 1A700, parB(S278C)-linker-HaloTag(C61V,C262A)::specR, trpC2 | Gruber lab |
| BSG3398 | 1A700, parB(E279C)-linker-HaloTag(C61V,C262A)::specR, trpC2 | Gruber lab |
| BSG3945 | 1A700, smc-Pk3 ftsY::specR, ΔparA: SpnParB::kanR, trpC2 | Gruber lab |
| BSG3949 | 1A700, smc-Pk3 ftsY::specR, recoded ParB::camR, trpC2 | Gruber lab |
| BSG4050 | 1A700, parB(E111Q)::kanR, trpC2 | This work |
| BSG4051 | 1A700, parB(E111H)::kanR, trpC2 | This work |
| BSG4052 | 1A700, parB(E111A)::kanR, trpC2 | This work |
| BSG4053 | 1A700, parB(E78Q)::kanR, trpC2 | This work |
| BSG4054 | 1A700, parB(E78H)::kanR, trpC2 | This work |
| BSG4055 | 1A700, parB(E78A)::kanR, trpC2 | This work |
| BSG4056 | 1A700, smc-Pk3 ftsY::specR, parB(E111Q)::kanR, trpC2 | This work |
| BSG4059 | 1A700, smc-Pk3 ftsY::specR, parB(E78Q)::kanR, trpC2 | This work |
| BSG4113 | 1A700, parB(A102C)-linker-HaloTag(C61V,C262A)::specR, trpC2 | This work |
| BSG4114 | 1A700, parB(R105C)-linker-HaloTag(C61V,C262A)::specR, trpC2 | This work |
| BSG4115 | 1A700, parB(H133C)-linker-HaloTag(C61V,C262A)::specR, trpC2 | This work |
| BSG4116 | 1A700, parB(L134C)-linker-HaloTag(C61V,C262A)::specR, trpC2 | This work |
| BSG4117 | 1A700, parB(A102C, H133C)-linker-HaloTag(C61V,C262A)::specR, trpC2 | This work |
| BSG4118 | 1A700, parB(A102C, L134C)-linker-HaloTag(C61V,C262A)::specR, trpC2 | This work |
| BSG4119 | 1A700, parB(R105C, H133C)-linker-HaloTag(C61V,C262A)::specR, trpC2 | This work |
| BSG4120 | 1A700, parB(R105C, L134C)-linker-HaloTag(C61V,C262A)::specR, trpC2 | This work |
| BSG4140 | 1A700, parB(T22C, E78Q)-linker-HaloTag(C61V,C262A)::specR, trpC2 | This work |
| BSG4141 | 1A700, parB(T22C, E111Q)-linker-HaloTag(C61V,C262A)::specR, trpC2 | This work |
| BSG4156 | 1A700, parB(A102, H133C, S278C)-linker-HaloTag(C61V,C262A)::specR, trpC2 | This work |
| BSG4356 | 1A700, Δsmc ftsY::ermB, parB(E111Q)::kanR, trpC2 | This work |
| BSG4357 | 1A700, Δsmc ftsY::ermB, parB(E78Q)::kanR, trpC2 | This work |
| BSG4666 | 1A700, parB-mScarlet::KanR, ΔcgeD::hbsU-mTorquois::CAT, trpC2 | This work |
| BSG4667 | 1A700, parB(E111Q)-mScarlet::KanR, ΔcgeD::hbsU-mTorquois::CAT, trpC2 | This work |
| BSG4668 | 1A700, parB(E78Q)-mScarlet::KanR, ΔcgeD::hbsU-mTorquois::CAT, trpC2 | This work |
| BSG4710 | 1A700, parB(E78Q A102C H133C S278C)-linker-HaloTag(C61V,C262A)::specR, trpC2 | This work |
| BSG4714 | 1A700, parB(R105C, H133C, S278C)-linker-HaloTag(C61V,C262A)::specR, trpC2 | This work |
| BSG4716 | 1A700, parB(A102C E111Q H133C S278C)-linker-HaloTag(C61V,-C262A)::specR, trpC2 | This work |
| BSG4718 | 1A700, parB(T22C, E78Q, S278C)-linker-HaloTag(C61V,C262A)::specR, trpC2 | This work |
| BSG4719 | 1A700, parB(T22C, E111Q, S278C)-linker-HaloTag(C61V,C262A)::specR, trpC2 | This work |
| BSG4793 | 1A700, parB(E78Q,R105C,H133C,S278C)-linker-HaloTag(C61V,C262A)::specR, trpC2 | This work |
| BSG4795 | 1A700, parB(R105C,E111Q,H133C,S278C)-linker-HaloTag(C61V,-C262A)::specR, trpC2 | This work |
| BSG4775 | 1A700, smc-Pk3 ftsY::tetR, parB(A102C,H133C)-linker-HaloTag(C61V,-C262A)::specR, trpC2 | This work |
| BSG4776 | 1A700, smc-Pk3 ftsY::tetR, parB(R105C,H133C)-linker-HaloTag(C61V,-C262A)::specR, trpC2 | This work |
| BSG4777 | 1A700, smc-Pk3 ftsY::tetR, parB(R105C,L134C)-linker-HaloTag(C61V,-C262A)::specR, trpC2 | This work |
| BSG4778 | 1A700, smc-Pk3 ftsY::tetR, parB(A102C)-linker-HaloTag(C61V,C262A)::specR, trpC2 | This work |
| BSG4779 | 1A700, smc-Pk3 ftsY::tetR, parB(R105C)-linker-HaloTag(C61V,C262A)::specR, trpC2 | This work |
| BSG4780 | 1A700, smc-Pk3 ftsY::tetR, parB(H133C)-linker-HaloTag(C61V,C262A)::specR, trpC2 | This work |
| BSG4781 | 1A700, smc-Pk3 ftsY::tetR, parB(L134C)-linker-HaloTag(C61V,C262A)::specR, trpC2 | This work |
| BSG4815 | 1A700, Δ8-parS, parB(T22C, E111Q, mtparS)-linker-HaloTag(C61V,-C262A)::specR, smc::ermB, trpC2 | This work |
| BSG4816 | 1A700, Δ8-parS, parB(T22C, E78Q, mtparS)-linker-HaloTag(C61V,-C262A)::specR, smc::ermB, trpC2 | This work |
| BSG4852 | 1A700, Δ8-parS, parB(mtparS)-linker-HaloTag(C61V,C262A)::specR, smc::ermB, trpC2 | This work |

**Table S2.**
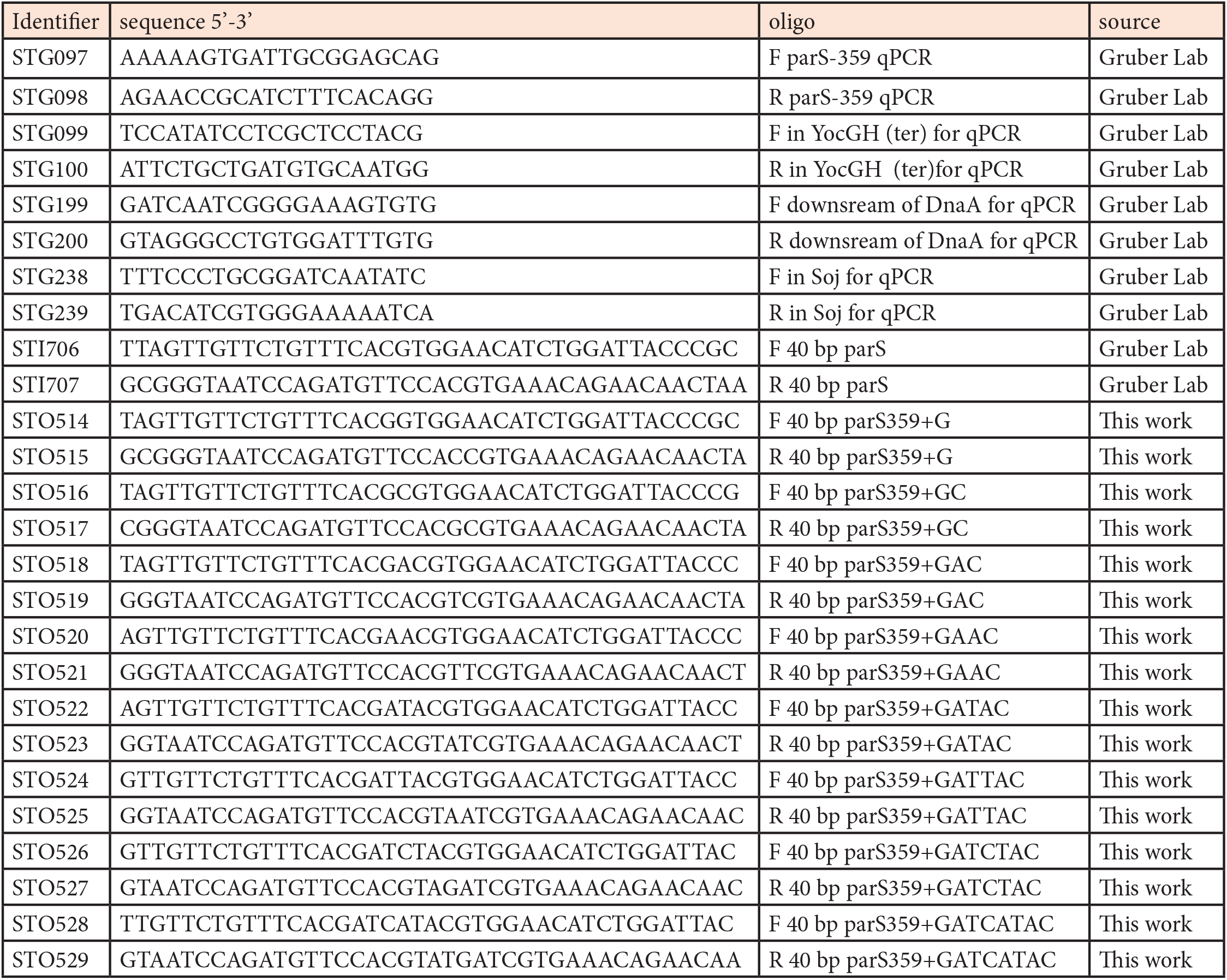
Primers used in this study.

